# Temporal niche partitioning through olfactory cell type evolution

**DOI:** 10.64898/2026.09.04.749392

**Authors:** Ambra Masuzzo, Asfa Sabrin Borbora, Rolando D. Moreira-Soto, Liliane Abuin, Michael Reichelt, Michele Marconcini, Steeve Cruchet, Markus Knaden, Daehan Lee, Richard Benton

## Abstract

Temporal niche partitioning enables species to use the same resource at different times, but the underlying mechanisms are unknown. We show that *Drosophila sechellia*, a specialist on *Morinda citrifolia* noni, exhibits narrow temporal preference for the most toxic, ripe fruit, reducing exposure to competition, parasitization and microbial infection that can occur on other stages. Chemical analysis highlighted ketones as signature odors of ripe noni. Comparative single-cell transcriptomic atlases revealed that *D. sechellia* has co-opted expression of a larval ketone receptor, Or45a, in an adult olfactory neuron population. The novel expression of *Or45a* can be ascribed solely to *cis*-regulatory changes and is sufficient to confer physiological sensitivity to ripe noni. Importantly, Or45a is required for *D. sechellia*’s ripe noni preference, acting redundantly with a second ketone receptor, Or85c/b, whose neuron population has expanded in *D. sechellia*. Our results provide an unprecedented link between cell type evolution and ecologically-advantageous, temporal niche partitioning.

## Introduction

Temporal niche partitioning is an evolutionary strategy that allows sympatric species to co-exist in the same geographical region and use the same resources at different times (Carothers and Jaksić, 1984; Kronfeld-Schorl and Dayan, 2003). This phenomenon is thought to minimize competition for limited resources or space, and reduce predation risk for prey species. The majority of described examples of temporal partitioning are where species exhibit distinct diel variation in activity patterns (e.g., foraging, mating), peaking during the day, night or twilight periods (Chen et al., 2019; Lear et al., 2021; Nakabayashi et al., 2021; Nichols et al., 2025). Over longer timescales, species can co-exist in the same niche during different seasons, likely determined by the preference for distinct temperature and humidity conditions in the environment (Correa and Winemiller, 2014; Li et al., 2026; Mena et al., 2025).

Between these extremes of time distinction by species – over 24 hours and over several months – ecological niches themselves are changing over days to weeks. For example, many plant-based resources for animals, including fruits and flowers, develop, mature and then decay, which is accompanied by alterations in chemical composition, nutritional value, visual appearance and texture. However, there have been only fairly limited demonstrations of temporal niche partitioning over this timescale. For example, fig wasps use volatile chemical cues characteristic of different stages of fig development to oviposit at different times (Proffit et al., 2007); similarly, the fruit pest *Drosophila suzukii* uses diverse sensory cues to choose ripe fruit for egg-laying (Karageorgi et al., 2017; Little et al., 2020). In a different context, parasitoid wasps exploit different developmental stages of *Drosophila* larvae for infection (Zhang et al., 2026). In most cases, little or nothing is known about the specific cues defining temporal preference, the underlying sensory detection mechanisms often remain entirely unexplored, and the selective advantages not explicitly demonstrated.

An interesting model clade to examine temporal niche partitioning is the trio of closely-related drosophilids, *Drosophila melanogaster*, *Drosophila simulans* and *Drosophila sechellia* (Figure 1A). While *D. melanogaster* and *D. simulans* are cosmopolitan generalists, *D. sechellia* is endemic to the Seychelles islands, where it has evolved a lifestyle exclusively on the “noni” fruit of the tropical shrub *Morinda citrifolia* (Auer et al., 2021; Jones, 2005). Ripe noni (or an abundant fruit chemical, octanoic acid) is highly toxic for other drosophilids and more divergent insects (Legal et al., 1994; Legal and Plawecki, 1995). Thus, specialization on a toxic resource might alleviate competition and predation (Auer et al., 2021; Jones, 2005; Salazar-Jaramillo and Wertheim, 2021), for which *D. sechellia* is likely to be particularly prone given its low fecundity (Auer et al., 2021; Jones, 2005; R’Kha et al., 1997). Importantly, other drosophilid species and parasitoid wasps can be found on other stages of noni (Matute and Ayroles, 2014; Salazar-Jaramillo and Wertheim, 2021) suggesting that *D. sechellia* can only benefit from a restricted time window of noni ripening. Indeed, field studies revealed that *D. sechellia*, but not other insects, emerge from noni fruit samples collected at the “ripe” stage (Salazar-Jaramillo and Wertheim, 2021). However, such studies in nature are necessarily limited in sample sizes and the resolution of temporal staging of noni fruits.

**Figure 1.**
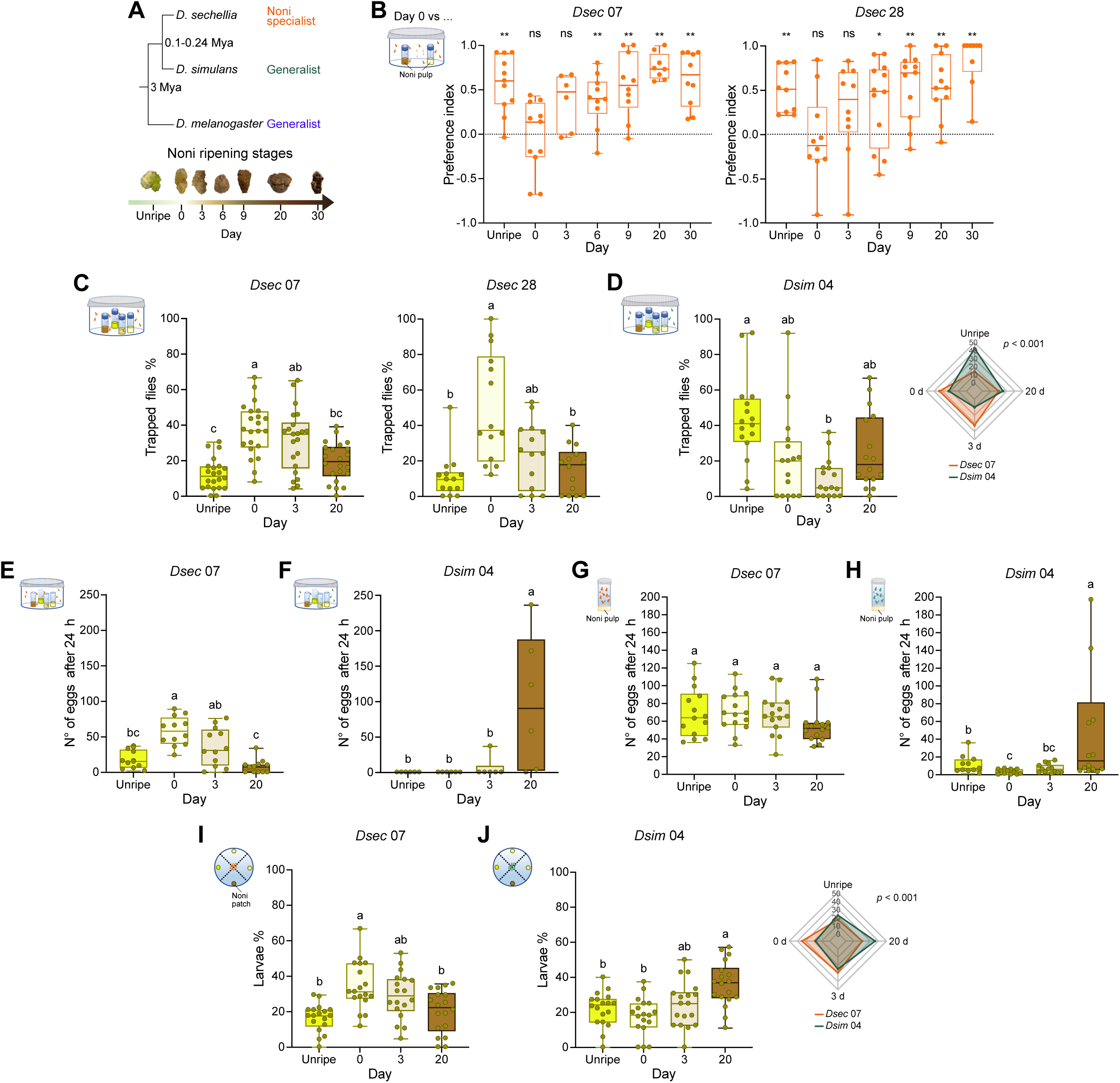
*D. sechellia* displays a narrow temporal window of host fruit preference. (A) Top: phylogeny of drosophilid species. Mya, millions of years ago. Bottom: all tested noni ripening stages: unripe, ripe (day 0, first day of ripeness), early overripe (days 3, 6 and 9), and late overripe (days 20 and 30). See Results and Methods for fruit staging details. (B) Behavioral responses of *D. sechellia* strains *Dsec* 07 and *Dsec* 28 (starved, mated females) in two-choice trap assays. Preference index for ripe (day 0) noni versus unripe, ripe (as control) or overripe stages is shown; *n* = 6-13 assays per strain. Positive values indicate preference for ripe noni. (C) Behavioral responses of *Dsec* 07 and *Dsec* 28 in multi-choice trap assays with four ripening stages. Box plots show the percentage of individuals trapped on each substrate; *n* = 22 (*Dsec* 07), *n* = 14 (*Dsec* 28). (D) Left: behavioral responses of *Dsim* 04 in multi-choice trap assays with four ripening stages. Box plots show the percentage of individuals trapped on each substrate. Right: radar plot comparing mean percentages of trapped flies per stage for *Dsim* 04 and *Dsec* 07 (control; d, day of ripeness).. *Dsim 04* (box and radar plots), *n* = 16; *Dsec 07* control (radar plot only), *n* = 11. (E,F) Oviposition site preference of *Dsec* 07 (E) and *Dsim* 04 (F) in multi-choice oviposition assays. Total egg counts on each substrate after 24 h are shown. For *Dsim* 04, replicates in which no eggs were laid were excluded prior to analysis; (E) *Dsec* 07, *n* = 12 (F) *Dsim* 04, *n* = 6. (G,H) No-choice oviposition assays of *Dsec* 07 (G) and *Dsim* 04 (H) on four noni stages. Raw egg counts on each substrate after 24 h are shown; (G) *Dsec* 07, *n* = 15 (H) *Dsim* 04, *n* = 14. (I) Behavioral responses of *Dsec* 07 larvae in chemotaxis assays with four noni ripening stages. Box plots show the percentage of larvae found on each substrate; *n* = 18. (J) Left: behavioral responses of *Dsim* 04 larvae in multi-choice chemotaxis assays with four ripening stages. Box plots show the percentage of larvae found on each substrate. Right: radar plot comparing mean percentages of larvae per stage for *Dsim* 04 and *Dsec* 07 (control; d, day of ripeness). *Dsim 04* (box and radar plots), *n* = 18; *Dsec 07* control (radar plot only), *n* = 18. In (B-J), box plots show the median and the first and third quartiles, with points representing individual biological replicates. *n* indicates the number of arenas (B-F), tubes (G,H) or plates (I,J). In (B) Wilcoxon signed-rank test. ns = non-significant, \**p* < 0.05, \*\**p* < 0.01, \*\*\**p* < 0.001. In (C,D) and (I,J), statistical analyses were performed on CLR-transformed values. Repeated measures one-way ANOVA with Tukey’s post-hoc test was performed for within-genotype comparisons with significant differences indicated by different letters (ɑ = 0.05). Mixed-design ANOVA with Genotype × Stage interaction was used for genotype comparisons (radar plots; *p* value for the difference in preference between genotypes is shown). In (E,F) Kruskal-Wallis test followed by Dunn’s post-hoc test with Bonferroni correction. Significant differences indicated by different letters (ɑ = 0.05). In (G,H) Negative binomial Generalized Linear Model (GLM) followed by Tukey’s post-hoc test. Significant differences indicated by different letters (ɑ = 0.05).

Here, we combine semi-natural behavioral assays, fruit volatile profiling, comparative single-cell transcriptomics, neurophysiology and genetic analysis to investigate the precision, molecular basis and evolution of temporal host specialization in *D. sechellia*. Our findings reveal how evolutionary changes in olfactory coding can refine host specialization from recognizing which resource to exploit to determining when to exploit it.

## Results

### *D. sechellia* displays a narrow temporal window of host fruit preference

We first standardized the noni ripening process using fruits harvested from greenhouse-grown plants (Figure 1A; see Methods). We defined seven ripening stages: an unripe stage (yellow-green fruit, 2-3 days before ripening), a ripe stage (day 0; soft texture, yellow-gray coloration, strongly pungent odor), and five overripe stages (days 3, 6, 9, 20 and 30, relative to the ripe stage). To determine *D. sechellia*’s temporal preference of noni, we performed two-choice trap assays; this paradigm examines only the influence of olfaction, which is likely to be the initial sensory modality for fruit choice (vision appears to be unimportant (Alvarez-Ocana et al., 2023)). Both tested *D. sechellia* strains strongly preferred ripe noni (day 0) over all other stages, except for early overripe (day 3) fruit (Figure 1B). These observations suggest that *D. sechellia* uses a narrow temporal window of its noni niche. For subsequent experiments, we selected four representative stages: unripe, ripe (day 0), early overripe (day 3) and late overripe (day 20).

We next established a more complex odor environment by performing multi-choice trap assays in which all four stages were presented simultaneously. *D. sechellia* robustly preferred day 0 and day 3 fruits (Figure 1C), while *D. simulans* and *D. melanogaster* showed no consistent preference for any stage (Figure 1D, Figure S1A). The preference of *D. sechellia* for day 0 and day 3 noni was robust across internal states (Figure S1B-D).

When we narrowed the temporal window to span days 0-3 of ripening, *D. sechellia* no longer showed a preference for day 0, suggesting that the olfactory cues sufficient to drive ripe stage attraction are present across most of this period (Figure S1E). We also tested *D. sechellia* preference in the multi-choice assay in the absence of the day 0 stage. In this case, flies preferred day 3 noni over both unripe and day 20 stages (Figure S1F), further supporting the idea that the key chemical cues are maintained through day 3.

We next asked whether the temporal preference for ripe noni extends to oviposition. This behavior involves direct contact with the substrate, resulting in taste and textural cues becoming accessible in addition to odors, raising the question of whether stage preference is maintained or modified. *D. sechellia* preferentially oviposited on day 0 and day 3 noni, consistent with its olfactory preference for these stages (Figure 1E). By contrast, *D. simulans* exclusively laid eggs on day 20 noni (Figure 1F). To assess whether noni-ripening stage affects fecundity independently of preference, we performed no-choice oviposition assays in which females were confined to a single stage (Figure 1G-H). *D. sechellia* laid comparable numbers of eggs across all stages, indicating that fecundity is independent of ripening stage preference. *D. simulans* females again laid eggs principally on day 20 stage.

Finally, we assessed whether a similar preference is exhibited by larvae, given noni fruits can ripen heterogeneously, exposing foraging larvae to multiple ripening stages within a single fruit. *D. sechellia* larvae preferred day 0 and day 3 noni (Figure 1I), similar to adults, whereas *D. simulans* larvae slightly preferred day 20 stage (Figure 1J).

Together, these results demonstrate that *D. sechellia* has evolved a temporally precise host preference restricted to a narrow ripening window. This preference is species-specific, and consistent across different internal states, developmental stages and behavioral paradigms.

### Ecological consequences of *D. sechellia*’s noni ripe stage preference

We next investigated the ecological relevance of *D. sechellia*’s temporal niche preference. One hypothesis is that *D. sechellia* selects a specific ripening stage to minimize exposure to other species that might reduce its reproductive fitness. We first studied the wasp *Leptopilina boulardi*, which parasitizes drosophilid larvae (Schlenke et al., 2007) and has been found on noni in the Seychelles (Salazar-Jaramillo and Wertheim, 2021). Survival assays exposing *L. boulardi* to the four ripening stages revealed that day 0 and day 3 noni were highly lethal, with the vast majority of wasps dying within two hours, while unripe and day 20 stages were barely toxic (Figure 2A). The susceptibility of this wasp to noni exposure – which appears to reflect sensitivity to the main noni toxin octanoic acid (Figure S2A) (Legal et al., 1994) – is thus highest at the preferred stages of *D. sechellia*. Examining canonical anti-parasitoid defenses, we found that *D. sechellia* exhibits egg retention in the presence of wasps (as in *D. melanogaster* (Lefevre et al., 2012; Lynch et al., 2016)) (Figure S2B), but showed reduced larval avoidance of parasitoid chemical cues (Ebrahim et al., 2015) (Figure S2C) and, as previously reported (Salazar-Jaramillo et al., 2014), larvae failed to encapsulate wasp eggs (Figure S2D). Noni toxicity might have relieved selective pressures to maintain such defenses.

**Figure 2.**
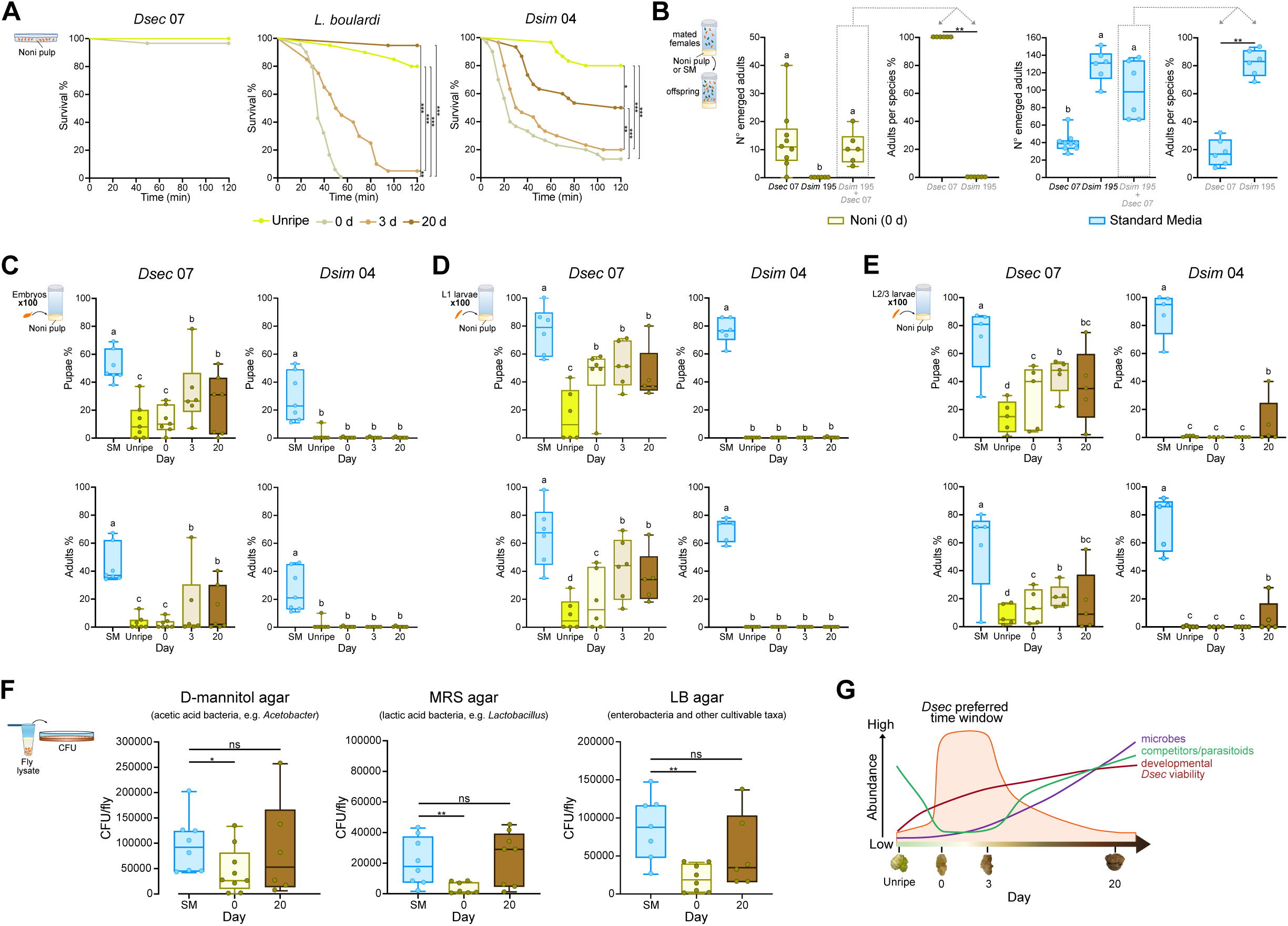
*D. sechellia* of noni ripe stage preference has fitness trade-offs. (A) Survivorship of *D. sechellia* (*Dsec* 07), *L. boulardi* (strain G486) and *D. simulans* (*Dsim* 04) adult females in plate assays on four noni stages. d, day of ripeness. Survival curves represent pooled data from all experiments; *n* = 20-30 animals per species. (B) Competition assays between *Dsec* 07 and *Dsim* 195 on ripe noni or standard media (SM). Mated females oviposited alone (10 females) or together (5 per species) for 24 h, and emerged adults were counted (schematic, left). For each substrate: number of emerged adults per condition. The grey dashed box indicates the mixed-species condition from which the species proportions in the adjacent graph are shown; *n* = 6-9 per condition. (C) Viability of *Dsec* 07 and *Dsim* 04 embryos on SM and four noni stages. Top: Percentage of pupae. Bottom: Percentage of adults; *n* = 6-7 per species. (D) Viability of *Dsec* 07 and *Dsim* 04 L1 larvae on SM and four noni stages. Top: Percentage of pupae. Bottom: Percentage of adults; *n* = 6 per species. (E) Viability of *Dsec* 07 and *Dsim* 04 L2/3 larvae on SM and four noni stages. Top: Percentage of pupae. Bottom: Percentage of adults; *n* = 4-5 per species. (F) Colony-forming unit (CFU) per fly from *Dsec* 07 reared on SM, ripe (day 0) or late overripe (day 20) noni, plated on D-mannitol, MRS or LB agar (see Methods); *n* = 6-9 per condition. (G) Model of the ecological trade-offs across noni ripening. Competitor and parasitoid survival and microbial load are low across the ripe to early overripe, while *D. sechellia* developmental viability increases with ripening. The shaded day 0-day 3 window marks the preferred noni stage for *D. sechellia*, where threats are minimal, yet viability remains sufficient to support offspring. In (A), Log-rank tests with Holm correction. \**p* < 0.05, \*\**p* < 0.01, \*\*\**p* < 0.001. In (B-F), box plots show the median and the first and third quartiles, with points representing biological replicates (*n*). In (B), for the graphs showing number of emerged adults: Kruskal-Wallis test followed by Dunn’s post-hoc test with Bonferroni correction. Significant differences indicated by different letters (ɑ = 0.05). For the graphs showing percentages of emerged flies per species: Mann-Whitney test. \*\**p* < 0.01. In (C-E) Binomial GLM with Firth’s correction and Tukey’s post-hoc test. Significant differences indicated by different letters (ɑ = 0.05). In (F) Mann-Whitney test. ns = non-significant, \**p* < 0.05, \*\**p* < 0.01.

We next focused on competition with *D. simulans*, which co-occurs with *D. sechellia* in the Seychelles (Matute and Ayroles, 2014). Like wasps, *D. simulans* was most sensitive to day 0 and day 3stages (Figure 2A). To test whether noni protects *D. sechellia* from this competitor, we reared the two species alone or together, on standard medium or day 0 noni (Figure 2B). As expected, no *D. simulans* offspring emerged on day 0 noni (Figure 2B). By contrast, on standard medium, *D. simulans* produced significantly more offspring than *D. sechellia*, consistent with the low fecundity of the latter (R’Kha et al., 1997). Moreover, when the two species were reared together, *D. sechellia* was strongly outnumbered by *D. simulans* (∼15% versus ∼85% of emerged adults), suggesting that, in the absence of noni, *D. sechellia* would be rapidly outcompeted. Thus, host-fruit toxicity might buffer *D. sechellia* against competitive exclusion by *D. simulans*, offsetting its intrinsic fecundity disadvantage.

To investigate further reproductive fitness effects, we compared the viability on noni of both species from embryos, first-instar, and second-/early third-instar larvae through pupation to adulthood (Figure 2C-E). Pupal and adult counts were highest on standard media for both species, indicating that noni is a suboptimal and/or toxic developmental diet for both. Surprisingly, viability of *D. sechellia* on noni was highest when it initiates on the two overripe stages (the substrate presumably changes over the period of animal development), suggesting that while *D. sechellia* adults strongly prefer day 0, embryonic and larval development is better supported by more advanced ripening. *D. simulans* produced no (or very few) pupae or adults on any noni substrate when starting from any developmental stage (Figure 2C-E), confirming the potent barrier that noni toxicity poses to *D. simulans* development.

Lastly, we asked how noni ripening stage shapes *D. sechellia*’s bacterial exposure by quantifying the total culturable bacterial load of flies reared on day 0 and day 20 noni – the latter assumed to be microbe-enriched due to its decomposing state – using a panel of selective and general media (Figure 2F). Rearing *D. sechellia* on day 0 noni lowered bacterial load across all groups, whereas day 20 noni exhibited similar levels to those of flies cultured on standard medium. The strong bacterial suppression at the day 0 stage – which is consistent with the low microbiome complexity of *D. sechellia* (Chandler et al., 2011; Heys et al., 2021) – might limit *D. sechellia*’s encounters with pathogenic microbes.

These results indicate that temporal preference for day 0 noni by *D. sechellia* is a trade-off: this ripening stage is the most toxic to other species, relieving potential competition, parasitization and infection, but also leads to lower viability of *D. sechellia*’s progeny compared to that on both overripe stages (Figure 2G).

### Changes in the noni odor bouquet during ripening

To understand the sensory basis of *D. sechellia*’s temporal host preference, we first characterized the volatile odor profile of noni fruit across ripening stages by solid-phase microextraction (SPME) coupled to gas chromatography-mass spectrometry (GC-MS), using behaviorally-validated fruit samples (Figure 3A-B). Each ripening stage displayed a distinctive profile of chemicals (Figure 3C). The noni bouquet is dominated by diverse esters enriched at one or a subset of ripening stages: short-chain esters such as pentyl and hexyl hexanoate were abundant only at the unripe stage, *4*-pentenyl propionate and *4*-pentenyl butyrate at the day 0 and day 3 stages, while the day 20 stage showed greater chemical diversity, likely reflecting advanced ripening and microbial fermentation (Mansourian and Stensmyr, 2015; Zhou et al., 2024).

**Figure 3.**
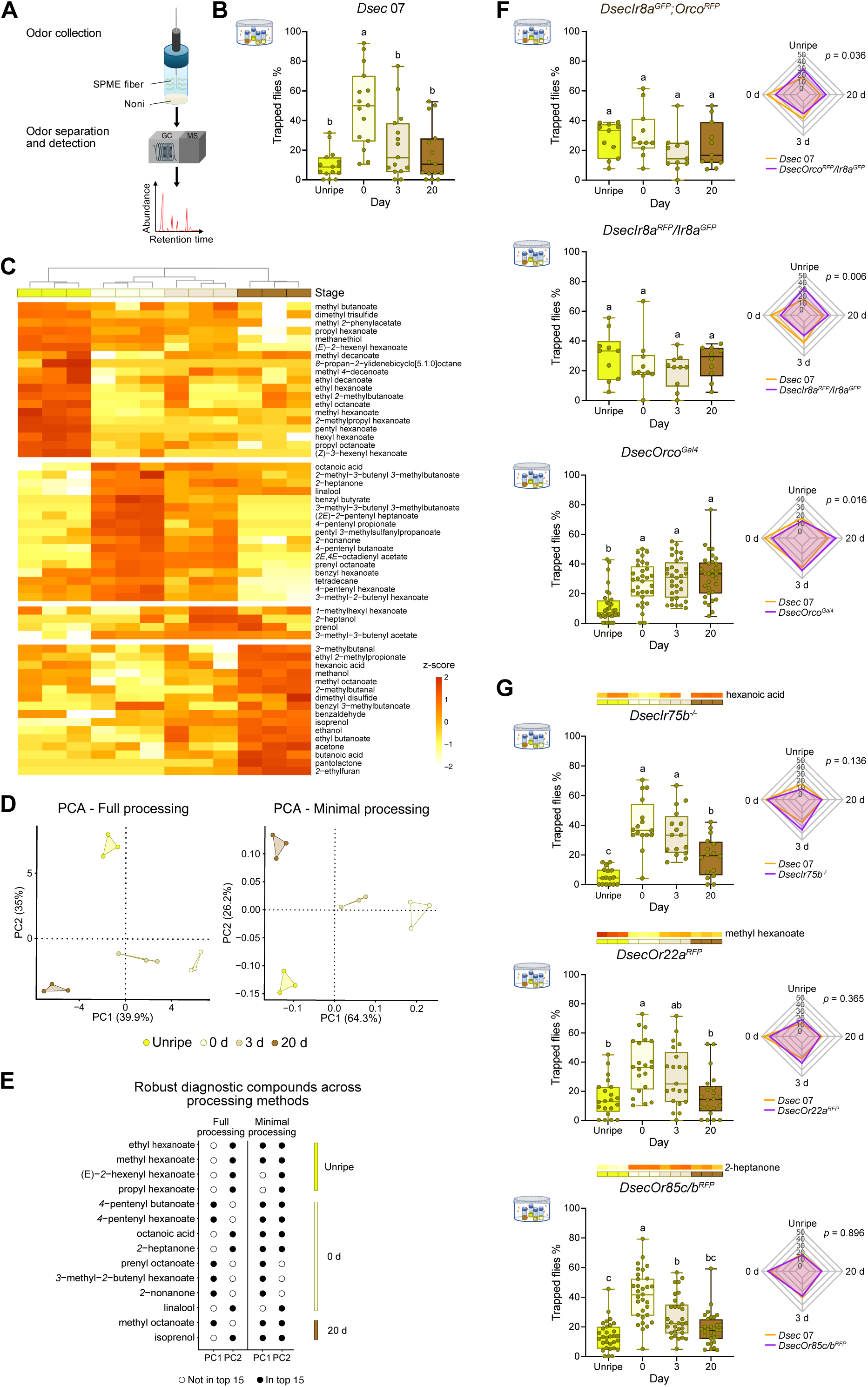
Changes in the noni odor bouquet during ripening. (A) Gas chromatography-mass spectrometry (GC-MS) experimental setup. Compounds were collected from the headspace of noni pulp at four ripening stages using solid-phase microextraction (SPME) fibers and separated by GC-MS. Three fruits per stage were analyzed and a total of 56 compounds were identified (see Table S7). (B) Behavioral responses of *Dsec* 07 in multi-choice trap assays with four ripening stages, using the same fruit samples as those used for GC-MS analysis. Box plots show the percentage of individuals trapped on each substrate; *n* = 15. (C) Hierarchical cluster analysis dendrogram of the 12 fruit samples (color-coded as in (B)), computed on autoscaled data using Euclidean distance and Ward’s linkage method. The heatmap below shows the relative abundance of all 56 detected volatile compounds across ripening stages. White gaps separate volatiles grouped by the ripening stage at which they are most abundant. Color scale represents *z*-scores calculated per compound across all samples. (D) Left: PCA score plot of fully processed data showing the separation of fruit samples by ripening stage based on their volatile profile. To give equal analytical weight to all detected compounds, peak intensities were normalized, log-transformed and autoscaled prior to PCA (see Methods). Right: Principal Component Analysis (PCA) score plot of minimally processed data showing the separation of fruit samples by ripening stage based on their volatile profile. Peak intensities were only normalized sample-wise, without further transformation or scaling, reflecting the absolute abundance of compounds (see Methods); d, day of ripeness. (E) Volatile compounds consistently ranking among the top 15 by absolute loading value across at least one PC (PC1 or PC2) in either the minimally or fully processed data. Black points indicate that the compound is within the top 15 for that PC; open points indicate it is not. (F, G) For each panel, the left shows behavioral responses of *Dsec* mutants in multi-choice trap assays with four ripening stages, and the right shows a radar plot comparing mean percentages of trapped flies per stage between *Dsec* 07 (control) and the indicated mutant genotype. Box plots show the percentage of individuals on each substrate. (F) *DsecOrco^RFP^/Ir8a^GFP^* double mutant (box and radar plots), *n* = 11; *Dsec* 07 (radar plot only), *n* = 19. *DsecIr8a^RFP^/Ir8a^GFP^* mutant (box and radar plots), *n* = 10; *Dsec* 07 (radar plot only), *n* = 14. *DsecOrco^Gal4^* mutant (box and radar plots), *n* = 32; *Dsec* 07 (radar plot only), *n* = 30. (G) *DsecIr75b^-/-^* mutant, (box and radar plots), *n* = 16; *Dsec* 07 (radar plot only), *n* = 17. *DsecOr22a^RFP^* mutant, (box and radar plots), *n* = 21; *Dsec* 07 (radar plot only), *n* = 19.. *DsecOr85c/b^RFP^* mutant(box and radar plots), *n* = 31; *Dsec* 07 (radar plot only), *n* = 51. In (B) and (F,G), box plots show the median and the first and third quartiles, with points representing biological replicates (arenas, *n*). Statistical analyses were performed on CLR-transformed values. Repeated measures one-way ANOVA with Tukey’s post-hoc test was performed for within-genotype comparisons, with significant differences indicated by different letters (α = 0.05). Mixed-design ANOVA with Genotype × Stage interaction was used for genotype comparisons (radar plots; *p* value for the difference in preference between genotypes is shown).

Principal component (PC) analysis confirmed tight clustering by ripening stage (Figure 3D). PC1 captured the main axis of chemical change across ripening, with day 0-stage compounds driving the positive loadings, while PC2 separated the unripe stage from all others, reflecting a distinct set of unripe volatiles that decline sharply upon ripening (Figure 3C-D and Figure S3A). This clustering was robust to the choice of data-processing approach (see Methods; Figure 3D and Figure S3B).

To identify the compounds most robustly diagnostic of each stage, we compared loadings across analyses and focused on those consistently appearing among the top 15 contributors (Figure 3E and Figure S3). The day 0 stage was defined by a core set including octanoic acid, linalool, four esters (*4*-pentenyl hexanoate, *4*-pentenyl butanoate, prenyl octanoate, and *3*-methyl-*2*-butenyl hexanoate) and the ketones *2*-heptanone and *2*-nonanone. No compounds were uniquely associated with the day 3 stage, which broadly resembled the day 0 stage but with lower chemical abundance, consistent with *D. sechellia’*s inability to reliably discriminate between these two stages. Several of these compounds – including methyl hexanoate and *2*-heptanone – have previously been associated with *D. sechellia*’s long-range attraction to noni (Auer et al., 2020; Dekker et al., 2006; Ibba et al., 2010).

Given that stage preference extends to oviposition and larval behavior, where non-volatile cues become accessible, we asked whether sugar and amino acid composition varies with ripening stage using liquid chromatography-mass spectrometry (Figure S4). Stage separation based on sugar content was less distinct than for volatiles, with the two overripe stages showing similarly elevated sugar abundances and no sugar particularly enriched at the day 0 stage (Figure S4A-B). Free amino acid profiles yielded stage-specific cluster separation, but similarly showed no set of amino acids uniquely enriched at the day 0 stage (Figure S4C-D). Thus, among the chemical properties of noni examined, the volatile compounds most distinctively mark the ripe stage.

### Noni-attraction receptors do not explain ripe-stage fruit discrimination

To dissect the sensory basis of *D. sechellia*’s ripe stage preference, we tested a panel of olfactory receptor mutants. Near-anosmic *D. sechellia* lacking the broadly-expressed olfactory co-receptors Ir8a (Abuin et al., 2011) and Orco (Benton et al., 2006; Larsson et al., 2004) showed no behavioral discrimination between fruit stages. The contributions of the Ionotropic receptor (Ir)-dependent and Odorant receptor (Or)-dependent pathways were not equivalent: mutants lacking only *Ir8a* also showed no stage discrimination, while *Orco* mutants retained lower attraction to unripe noni (Figure 3F).

We next sought the ligand-specific (“tuning”) receptor(s) that act with these co-receptors (Figure 3G). The best-characterized tuning Ir in *D. sechellia* is Ir75b, which has evolved novel sensitivity to hexanoic acid and promotes short-range attraction and oviposition (Alvarez-Ocana et al., 2023; Prieto-Godino et al., 2017). However, hexanoic acid was detected at the lowest levels at the day 0 stage and *Ir75b* mutants still discriminate noni stage similarly to wild type animals (Figure 3G), suggesting that this sensory pathway is not important for ripe fruit preference. The best-characterized *D. sechellia* tuning Or is Or22a, which has evolved heightened sensitivity to methyl hexanoate (compared to *D. melanogaster*) and mediates long-range attraction to noni (Auer et al., 2020; Dekker et al., 2006). Unexpectedly, methyl hexanoate was most abundant in unripe fruit extracts, and *Or22a* mutants retained preference for the day 0 stage (Figure 3G), arguing against a central role for this receptor in ripening stage discrimination. Another candidate receptor is Or85c/b (representing two very closely-related, functionally-similar paralogs, Or85c and Or85b (Auer et al., 2020)), which detects *2*-heptanone, one of the two ketones robustly enriched at the day 0 stage (Figure 3C,E). However, *Or85c/b* mutants also retained ripe preference (Figure 3G). Together, these results suggested that stage discrimination relies on combinatorial coding by multiple receptors and/or on olfactory pathways not yet characterized in *D. sechellia*.

### Identification of novel *D. sechellia*-specific changes in olfactory pathways

To identify *D. sechellia* olfactory pathways that contribute to temporal stage discrimination, we took an unbiased comparative approach, by surveying the molecular and cellular composition of the entire antennal olfactory system of *D. sechellia*, *D. simulans* and *D. melanogaster* through single-nucleus RNA-sequencing (snRNA-seq) (Figure 4A). Within the atlases of OSNs of the three species, homologous populations of OSNs were readily identified (Figure 4B-C and Figure S5A-B). Comparison of the frequency of each neuron type revealed global conservation in the representation of both Or and Ir neuron populations (Figure 4D and Figure S5C). However, *D. sechellia* exhibits several differences compared to the other species. Several neuron populations have decreased representation in *D. sechellia*, including those expressing *Or47a*, *Or13a* and *Or9a* (Figure 4D), consistent with earlier *in situ* analyses of these populations (Auer et al., 2020; Takagi et al., 2024). In *D. melanogaster*, several of these pathways detect odors associated with yeast alcoholic fermentation (Munch and Galizia, 2016; Schlegel et al., 2021), raising the possibility that their decreased representation in *D. sechellia* reflects their dispensability for detecting ripe noni fruit.

**Figure 4.**
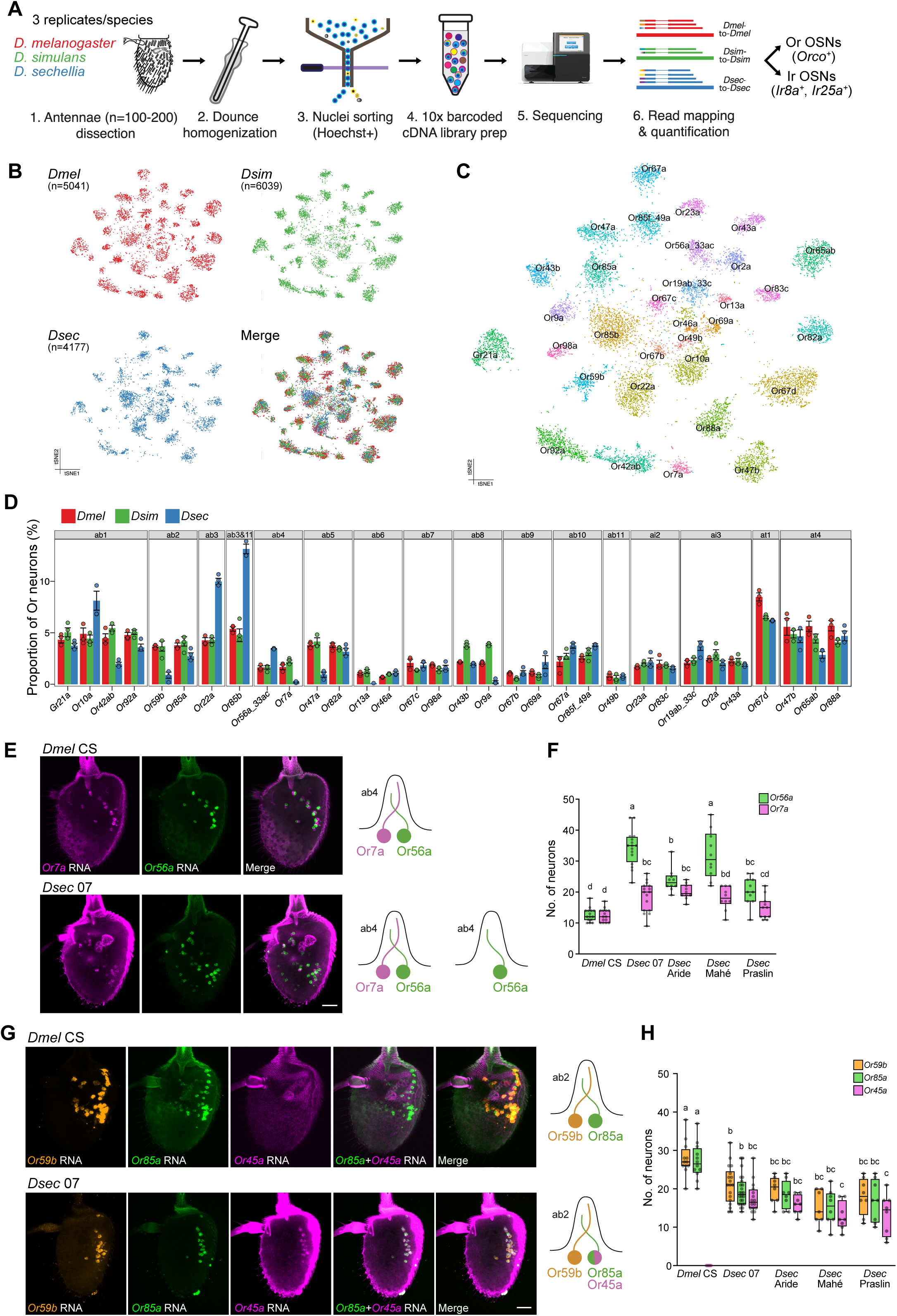
Comparative antennal snRNA-seq to identify *D. sechellia*-specific changes in olfactory pathways. (A) Schematic of the comparative antennal snRNA-seq workflow. (B) tSNE plots of *D. melanogaster* (*Dmel* CS), *D. simulans* (*Dsim* 04) and *D. sechellia* (*Dsec* 07) antennal Or subsystem neurons from an integrated dataset after RPCA integration; the number of cells in each atlas is indicated (*n*). In the bottom right plot, all cells from the three species are merged. (C) tSNE plot of the merged dataset from (B) in which cluster identity (neuron type) is annotated. (D) Bar plot illustrating Or subsystem neuron type frequency comparisons across species, organized by sensilla: ab, antennal basiconic; at, antennal trichoid; ai, antennal intermediate (Or85b neurons are housed in both ab3 and ab11 sensilla (Benton et al., 2025)). Each point corresponds to one of the three biological replicates. (E) Left: RNA FISH for *Or7a* and *Or56a* in whole-mount antennae of *Dmel* CS and *Dsec* 07. Scale bar, 25 µm. Right: schematic of inferred ab4 sensillar organization. (F) Quantification of ab4 neurons in *Dmel* CS and *Dsec* 07 and recently wild-caught strains of *D. sechellia* (Shahandeh et al., 2026)). *n* = 7-24 sensilla per neuron type. (G) Left: RNA FISH for *Or59b*, *Or85a* and *Or45a* in whole-mount antennae of *Dmel* CS and *Dsec* 07. Scale bar, 25 µm. Right: schematic of inferred ab2 sensillar organization. (H) Quantification of ab2 neurons in the indicated species and strains. *n* = 7-24 sensilla per neuron type. In (F) and (H), data are presented as box plots; center line, median; whiskers, minimum to maximum values; one-way ANOVA with Tukey’s post-hoc test was performed with significant differences indicated by different letters (ɑ = 0.05).

Other populations displayed elevated representations in *D. sechellia*, suggestive of increased ecological relevance for *D. sechellia*. These include the Or22a and Or85c/b neuron populations (co-housed in antennal basiconic 3 (ab3) sensilla), and the Ir75b population (in antennal coeloconic 3 I (ac3I)), again concordant with *in situ* analyses (Auer et al., 2020; Dekker et al., 2006; Prieto-Godino et al., 2017; Takagi et al., 2024). We also observed an increased representation of Or56a neurons – which detect geosmin, a volatile chemical that signals the presence of harmful microbes (Stensmyr et al., 2012) – and decreased representation of the co-housed Or7a neurons in ab4 (Figure 4D). We confirmed an increased population size of *D. sechellia Or56a* neurons by HCR-FISH both in our laboratory strains and in more recently wild-caught *D. sechellia* lines (Shahandeh et al., 2026) (Figure 4E-F). While we could detect Or7a neurons in *D. sechellia*, the imbalanced representation might in part be explained by our observation of many Or56a neurons that were unpaired with Or7a neurons, suggestive of an additional type of ab4 sensillum in which Or7a neurons are not specified (or die) during development (Figure 4E). We suggest that the increased sensory representation of geosmin-sensing neurons in *D. sechellia* helps this species avoid overripe, microbe-rich substrates.

### Co-option of the larval receptor Or45a in adult *D. sechellia*

None of the peripheral changes described above can explain the attraction of *D. sechellia* to day 0 noni. We therefore turned our attention to the ab1 and ab2 sensilla, for which earlier electrophysiological analyses suggested the occurrence of more substantial modifications to their functions in *D. sechellia* (Dekker et al., 2006; Stensmyr et al., 2003). As cross-species integration of snRNA-seq atlases can sometimes artefactually force different cell types into the same cluster, we first extracted from our integrated atlas all the clusters expressing known ab1 and ab2 receptors, and re-clustered these by species for both *D. melanogaster* and *D. sechellia* (Figure S5D-E).

In *D. melanogaster*, we could identify four well-defined cell clusters corresponding to the known ab1 neuron types (expressing *Or42b*, *Or92a*, *Gr21a* and *Or10a*) and two well-defined cell clusters corresponding to the known ab2 neuron types (expressing *Or59b* and *Or85a*). In *D. sechellia*, several differences were observed. For ab1, we could identify 3 neuron types (Or92a, Gr21a, Or10a), but Or42b neurons were completely absent (Figure S5D). Similar to the other diminished neuronal populations described above, this loss might reflect the lack of requirement for this sensory pathway which is important for detecting fermentation products in *D. melanogaster* (Semmelhack and Wang, 2009). For ab2 neurons, we identified one cell cluster expressing only *Or59b* and, more surprisingly, another cluster containing cells expressing *Or85a*, *Or59b* (albeit more sparsely than the other ab2 cluster) and a third receptor, *Or45a* (Figure S5D-E). In *D. melanogaster*, Or45a is a larval-specific receptor (Fishilevich et al., 2005; Kreher et al., 2005); although this gene’s expression in the *D. sechellia* antenna has been observed before through bulk RNA-seq (Bontonou et al., 2024; Shiao et al., 2015), the cell type in which it is expressed was unknown. The *D. sechellia*-specific expression of Or45a caught our attention as this is a sensitive receptor for *2*-nonanone and *2*-heptanone (Kreher et al., 2005; Mathew et al., 2013), which are odors characteristic of ripe noni (Figure 3C,E).

To validate the RNA-seq predictions, we performed RNA FISH for *Or85a*, *Or59b* and *Or45a* in *D. melanogaster* and *D. sechellia* (Figure 4G-H). In both species, the two ab2 neurons, *Or59b* and *Or85a* are paired, although there is a small reduction of ab2 sensilla numbers in *D. sechellia*. Most importantly, we observed co-expression of *Or45a* with *Or85a* in *D. sechellia* but not in *D. melanogaster* (Figure 4G-H).

### Or45a is essential for detecting noni odors in *D. sechellia*

To test the functional significance of the novel *Or45a* expression in *D. sechellia*, we characterized the electrophysiological responses of ab2 neurons. In *D. melanogaster*, neuronal spike traces from ab2 sensilla are characterized by two distinct amplitudes corresponding to the ab2A neuron (which expresses Or59b and responds to methyl acetate) and the ab2B neuron (expressing Or85a, which responds to ethyl-*3*-hydroxybutyrate). We readily detected similar ab2 sensilla in *D. sechellia*, with comparable responses of the ab2A neuron to methyl acetate as in *D. melanogaster* (Figure 5A), but much lower sensitivity of the ab2B neurons to ethyl-*3*-hydroxybutyrate (Figure 5B). Consistent with the co-expression of *Or45a* with *Or85a*, the ab2B neuron in *D. sechellia*, but not *D. melanogaster*, also responded sensitively to *2*-nonanone and *2*-heptanone (Figure 5C-D), two of the best ligands for this larval receptor in *D. melanogaster* (Kreher et al., 2005; Mathew et al., 2013).

**Figure 5.**
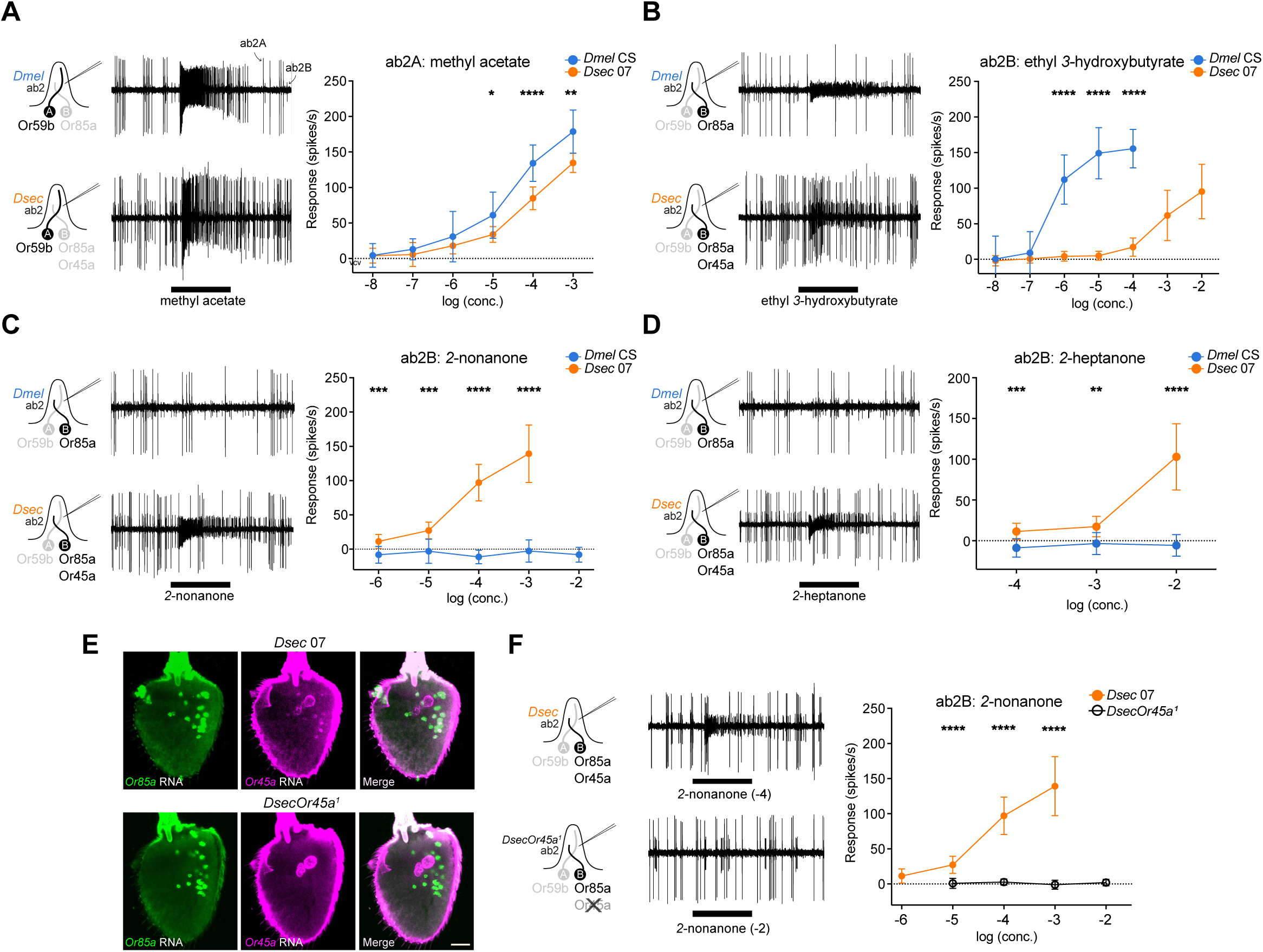
Or45a as a novel receptor for noni odors in *D. sechellia* antenna. (A-D) Left: representative traces of electrophysiological responses of ab2 sensilla neurons to the indicated odors. The black horizontal bar indicates the odor stimulus period (1 s). Right: quantification of dose-dependent ab2 neuron responses; *n =* 10-14 per species and odor concentration. (E) RNA FISH for *Or45a* and *Or85a* in whole-mount antennae of *Dsec* 07 and *DsecOr45a^1^* mutants. Scale bar, 25 µm. (F) Left: representative trace and right: quantification of ab2 neuron dose-response to the Or45a ligand *2*-nonanone in wild-type *Dsec* 07 and *DsecOr45a^1^* mutants; black horizontal bar, odor stimulus (1 s); *n* = 6-12 sensilla per strain and odor concentration. Data from *Dsec* 07 are the same as in (C). In (A-D) and (F), data are represented as mean ± SD; Shapiro-Wilk test for normal distribution, followed by unpaired two-tailed Student’s t-test comparison between neuronal responses at each concentration; \**p* < 0.05, \*\**p* < 0.01, \*\*\**p* < 0.001, \*\*\*\**p* < 0.0001.

In the course of these recordings, we noted the presence of a subclass of large basiconic sensilla in *D. sechellia* containing two neurons of nearly indistinguishable spike amplitude (Figure S6A). These sensilla did not respond to ligands of the other large basiconic sensilla, ab1 (CO_2_, which activates the Gr21a neuron) or ab3 (methyl hexanoate, which activates the Or22a neuron), but display identical responses to odors that activate neurons in the “typical” *D. sechellia* ab2 sensilla, including *2*-nonanone and *2*-heptanone (Figure S6A-D), suggesting they represent a distinct “atypical” class of ab2. The reason for the heterogeneity in neuronal spike amplitudes of typical and atypical ab2 sensilla in *D. sechellia* is unclear (see Discussion), but this phenotype – together with the changes in response profile of ab2B – might explain why ab2 sensilla were not clearly recognized in previous studies (Dekker et al., 2006; Stensmyr et al., 2003).

To determine whether the responses to *2*-nonanone in *D. sechellia* ab2 sensilla are due to the expression of Or45a (rather than altered tuning of Or85a) we generated an *Or45a* null mutant (Figure S6E). In these mutants, *Or45a* transcripts are no longer detected in *Or85a* neurons (Figure 5E), confirming loss of the receptor, but not the neuron in which it is normally expressed. Consistently, electrophysiological recordings from “typical” ab2 sensilla revealed the presence of two clear spike amplitudes, as in wild-type sensilla, but responses of ab2B to *2*-nonanone were completely absent (Figure 5F). Thus, co-option of Or45a in adult *D. sechellia* confers novel odor sensitivity to the Or85a neuron.

### Redundant physiological and behavioral coding of noni stage specificity

To test if Or45a contributes to noni stage discrimination, we first examined the electrophysiological responses of these neurons to fruit at different ripening stages. In *D. melanogaster*, the ab2B neuron displays minimal responses to noni fruit stimuli of any stage. By contrast, *D. sechellia* ab2B neurons respond to all four stages, but with the highest responses to the day 0 stage (Figure 6A). These responses are due to Or45a, as they are absent in *D. sechellia Or45a* mutants (Figure 6A). These results provide evidence that Or45a encodes noni ripening stage information in *D. sechellia*.

**Figure 6.**
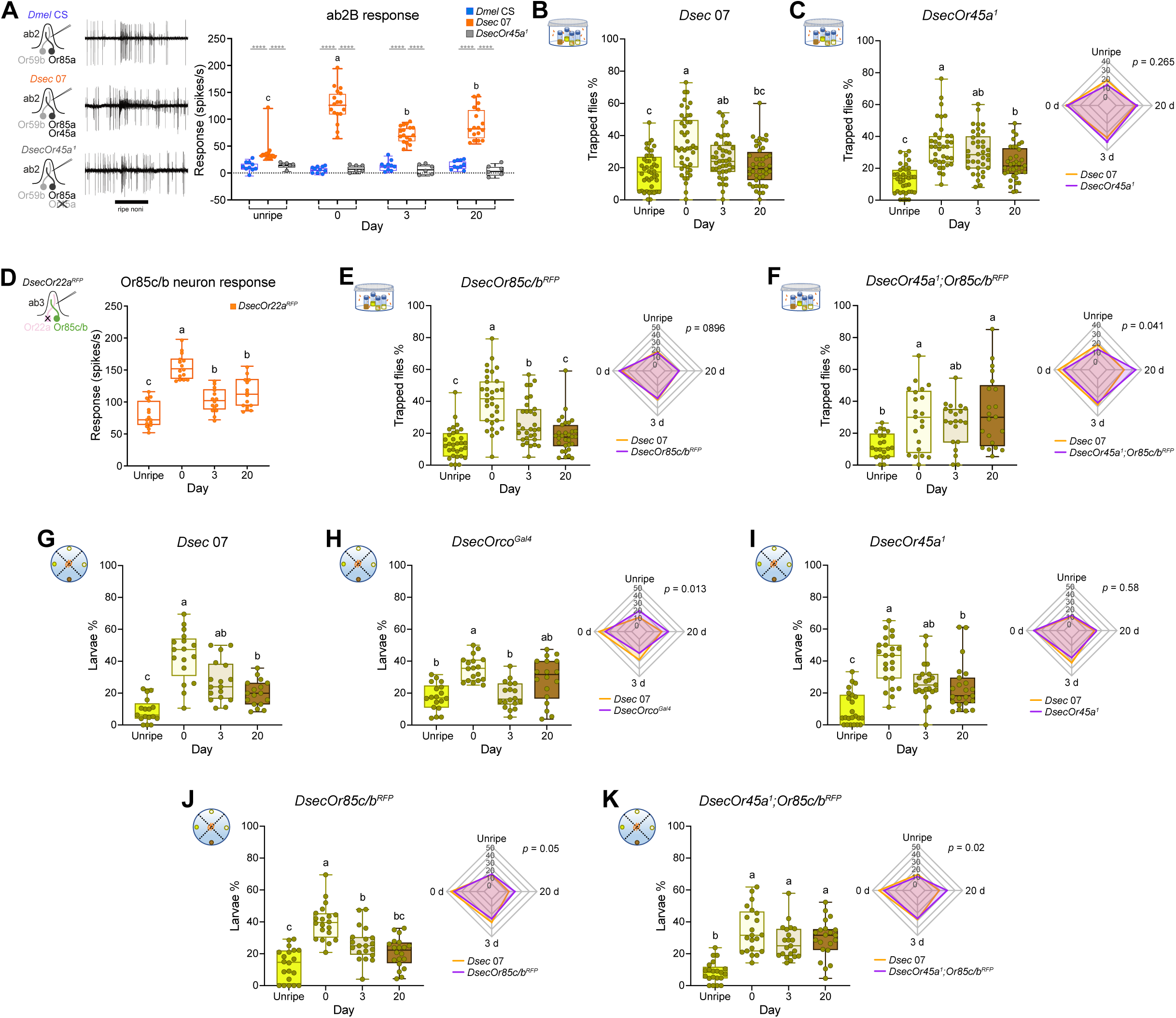
Unique and redundant physiological and behavioral coding of noni stage specificity. (A) Left: representative traces of electrophysiological responses of ab2 sensilla in *Dmel* CS*, Dsec* 07, and *DsecOr45a^1^*; black horizontal bar, odor stimulus (1 s). Right: quantification of ab2B neuron responses to noni fruit (50 mg) at different ripening stages; *n* = 6-17 sensilla per stage. (B) Behavioral responses of *Dsec* 07 in multi-choice trap assays with four ripening stages. Box plots show the percentage of individuals trapped on each substrate; *n* = 44. In (C-F), the left plot shows behavioral responses of *Dsec* mutants in multi-choice trap assays with four ripening stages, and on the right is a radar plot comparing mean percentages of trapped flies per stage between the indicated mutant genotype and *Dsec* 07 (control; d, day of ripeness). Box plots show the percentage of individuals on each substrate.(C) *DsecOr45a^1^* mutant (box and radar plots), *n* = 37; *Dsec* 07 (radar plot only), *n* = 44.. (E) *DsecOr85c/b^RFP^* mutant (box and radar plots), *n* = 31; *Dsec* 07 (radar plot only), *n* = 51. Data are replotted from Figure 3E. (F) *DsecOr85c/b^RFP^;Or45a^1^* double mutant, (box and radar plots), *n* = 21; *Dsec* 07 (radar plot only), *n* = 36. (D) Electrophysiological responses of Or85c/b neurons to noni fruit (10 mg) at different ripening stages; *n* = 15 sensilla per stage. The *DsecOr22a^RFP^* mutant background was used for these experiments to eliminate the responses of the Or22a neuron, which is co-housed with Or85c/b in the ab3 sensilla and responds to noni odors, resulting in pinching of the large spike amplitude, making it indistinguishable from the small spike of the Or85c/b neuron in the response window. (G) Behavioral responses of *Dsec* 07 larvae in multi-choice chemotaxis assays with four ripening stages. Box plots show the percentage of individuals found on each substrate; *n* = 17. (H-K) For each panel, the left shows behavioral responses of *Dsec* mutants in multi-choice chemotaxis assays with four ripening stages, and the right shows a radar plot comparing mean percentages of larvae per stage between the indicated mutant genotype and *Dsec* 07 (control; d, day of ripeness). Box plots show the percentage of larvae found on each substrate. (H) *DsecOrco^Gal4^* mutant, (box and radar plots), *n* = 18; *Dsec* 07 (radar plot only), *n* = 17. (I) *DsecOr45a^1^* mutant (box and radar plots), *n* = 23; *Dsec* 07 (radar plot only), *n* = 24. (J) *DsecOr85c/b^RFP^* mutant, (box and radar plots), *n* = 20; *Dsec* 07 (radar plot only), *n* = 18. (K) *DsecOr85c/b^RFP^;DsecOr45a^1^* double mutant, (box and radar plots), *n* = 21; *Dsec* 07 (radar plot only), *n* = 19*n* = 21. All box plots show the median and the first and third quartiles, with points representing biological replicates. In (B,C) and (E-K), n represents the number of arenas or plates. In (A) and (D), one-way ANOVA with Tukey’s post-hoc test for inter-stage comparisons, with significant differences indicated by different letters (α = 0.05). In (A), Mann-Whitney test for inter-genotypes comparison for each stage; \*\*\*\**p* < 0.0001. In (B,C) and (E-K), statistical analyses were performed on CLR-transformed values. Repeated measures one-way ANOVA with Tukey’s post-hoc test was performed for within-genotype comparisons, with significant differences indicated by different letters (α = 0.05). Mixed-design ANOVA with Genotype × Stage interaction was used for genotype comparisons (radar plots; *p* value for the difference in preference between genotypes is shown).

We next asked whether Or45a is required for behavioral discrimination of noni ripening stages. In multi-choice and two-choice trap assays, *Or45a* mutants display no differences in stage discrimination compared to wild-type *D. sechellia* (Figure 6B-C). Given the physiological evidence supporting Or45a’s candidacy, we hypothesized that this receptor might function partially redundantly with another olfactory pathway to define stage preference. A strong candidate for such a receptor is Or85c/b, expressed in the ab3B OSNs, whose neuronal population has expanded in *D. sechellia* (Figure 4D) (Takagi et al., 2024), and which is highly sensitive to *2*-heptanone (Auer et al., 2020; Ibba et al., 2010), a ripe noni-enriched ketone (Figure 3C,E). Supporting this possibility, *D. sechellia* Or85c/b neurons display the strongest electrophysiological responses to the odor bouquet of ripe noni (Figure 6D), similar to Or45a neurons.

As described above, *Or85c/b* mutants retained ripe stage preference (Figure 3G and Figure 6E). To test whether *Or45a* and *Or85c/b* act together in ripe stage discrimination, we generated flies lacking both of these receptors. Importantly, *Or45a;Or85c/b* double mutant combinations failed to discriminate the day 0 stage from overripe stages, resembling *Orco* mutants (the unripe stage is still discriminated by these mutants) (Figure 6F). These results indicate that Or45a and Or85c/b act redundantly as tuning receptors underlying ripe-stage preference in *D. sechellia*.

We next asked whether these olfactory pathways also underlie larval ripe stage preference. *Orco* mutant larvae failed to discriminate ripening stages (Figure 6G-H), confirming that *Or*-dependent olfaction is required for stage discrimination in these animals. While neither *Or45a* nor *Or85c/b* single mutant larvae showed impaired preference for day 0 noni (Figure 6I-J), the combined loss of these receptors abolished ripe stage preference (unripe noni was still avoided) (Figure 6K). Together, these results show that the same two olfactory receptors, Or45a and Or85c/b, act redundantly to mediate ripe-stage preference in both adults and larvae.

### *Cis*-regulatory changes underlie the novel expression of *Or45a* in *D. sechellia*

Finally, we sought to characterize the mechanisms underlying the novel adult expression of *Or45a* in *D. sechellia*. To examine the contribution of *cis*-regulatory evolution, we generated promoter-Gal4 driver transgenes containing ∼0.7 kb sequence upstream of the *Or45a* coding sequence from both *D. sechellia* and *D. melanogaster*. These were inserted into a common genomic location in *D. melanogaster* to avoid positional effects and eliminate the contribution of differences in the *trans* environment. *DmelOr45a-Gal4* drove expression of a *UAS-mCD8:GFP* effector transgene in some *Or85a*-negative antennal neurons (Figure 7A,E and Figure S7A), which presumably reflects a technical artefact (e.g., lack of inhibitory regulatory sequences) as *Or45a* is not expressed in adult *D. melanogaster* (Figure 5G). By contrast, *DsecOr45a-Gal4* robustly induced reporter expression in Or85a neurons, in addition to the same background expression (Figure 7B,E-F and Figure S7B). These results indicate that *cis*-regulatory changes within a short 5’ sequence of *D. sechellia Or45a* are sufficient to drive novel expression in the adult antenna.

**Figure 7.**
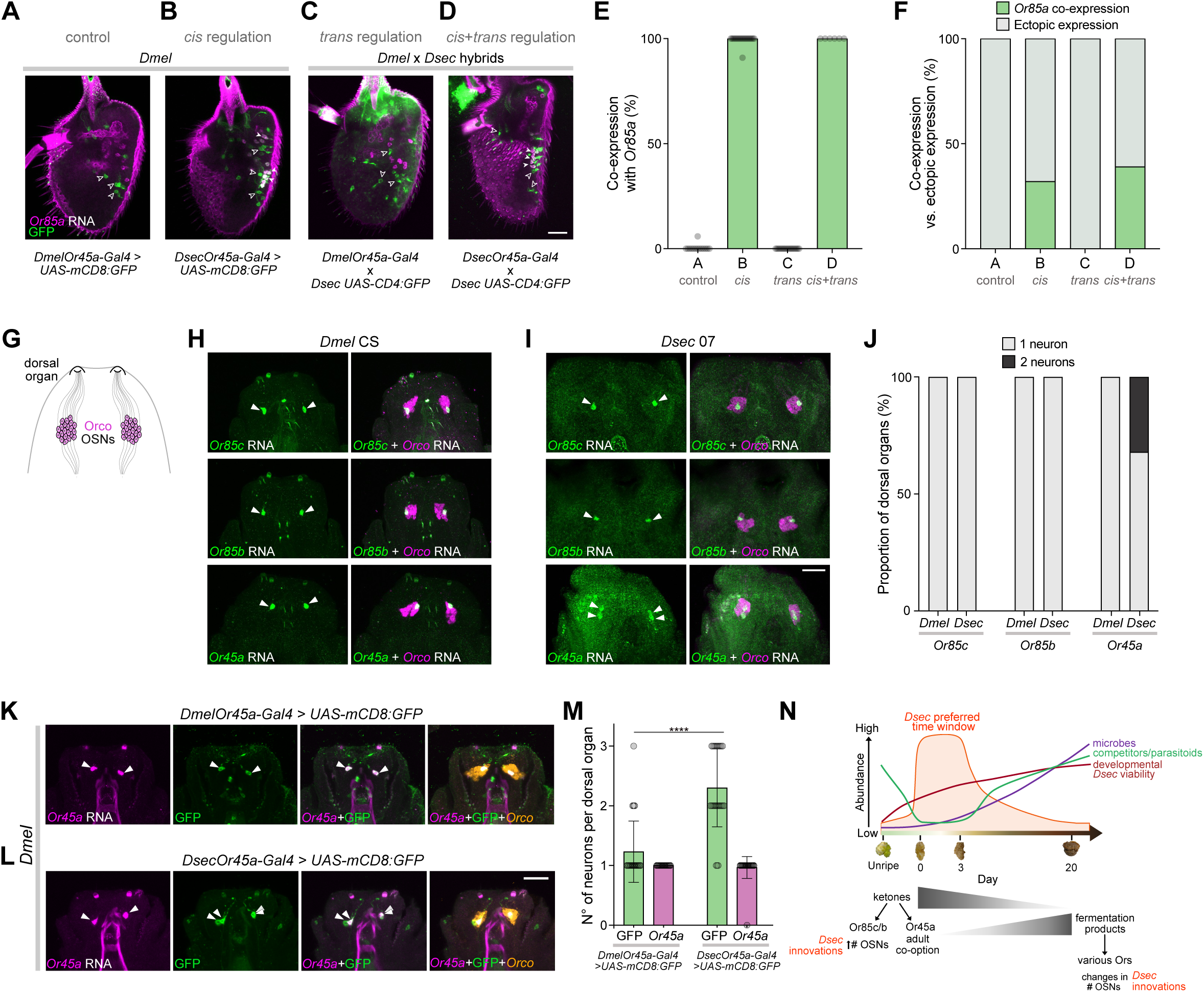
*Cis*-regulatory changes underlie the novel expression of *Or45a* in *D. sechellia*. (A-D) Immunofluorescence for GFP and RNA FISH for *Or85a* on antennae of *D. melanogaster* (A-B) or *D. melanogaster/D. sechellia* hybrids (C-D) of the indicated genotypes. Examples of ectopic expression of GFP are indicated with open arrowheads, and co-expression of GFP in *Or85a* neurons is indicated by closed arrowheads. Scale bar, 25 µm. (E) Percentage of *Or85a* neurons expressing GFP in the genotypes shown in (A-D); *n* = 6-20. (F) Percentage of neurons expressing GFP in *Or85a* neurons vs ectopic expression in the genotypes shown in (A-D); *n* = 6-20. (G) Schematic of the larval olfactory system. Cell bodies of 21 Orco-positive OSNs reside in the dorsal organ ganglion, with dendrites projecting to the peripheral dorsal organ. (H-I) RNA FISH for *Or85c*, *Or85b* or *Or45a*, together with *Orco*, in *D. melanogaster* (H) and *D. sechellia* (I) larval dorsal organs. Arrowheads indicate the cell bodies of GFP-labeled neurons in the dorsal organ ganglion. Scale bar, 25 µm. (J) Percentage of dorsal organs containing one or two neurons in (H) and (I); *n* = 18-34 dorsal organs from 9-18 larvae. (K-L) Immunofluorescence for GFP and RNA FISH for *Or45a* and *Orco* in larval dorsal organs of *D. melanogaster* larvae of the indicated genotypes. Arrowheads indicate the cell bodies of neurons in the dorsal organ ganglion. Scale bar, 25 µm. (M) Quantification of *Or45a*-positive and GFP-positive neurons in (J-K); *n* = 26-30 dorsal organs from 13-15 larvae. (N) Schematic adapted from (Figure 2G) with the addition of the sensory mechanisms underlying noni fruit stage discrimination. Data in (M) are presented as bar plots with mean ± SD; Mann-Whitney test; **** *p <* 0.0001.

To examine if there is any role of *trans*-regulation in *Or45a* expression, we made hybrids of *D. melanogaster* carrying *DmelOr45a-Gal4* (or *DsecOr45a-Gal4*) and *D. sechellia* bearing *UAS-CD4:GFP*. If there is a role of *trans*-regulation in inducing the adult expression of *Or45a*, the *D. melanogaster* promoter would be activated by transcription factors encoded within the *D. sechellia* half of the genome in the hybrid. However, the expression pattern of *DmelOr45a-Gal4* in the hybrid background did not differ from its expression in the *D. melanogaster* background (again, displaying only the same ectopic expression) (Figure 7C,E-F). As expected, *DsecOr45a-Gal4* induced reporter expression in Or85a neurons in the hybrids (Figure 7D,E-F). These results imply that *trans* regulation does not contribute to regulation of *Or45a* expression.

Given the requirement for Or45a and Or85c/b in noni stage preference in larvae, we also analyzed the expression of these receptors in the larval olfactory system, the dorsal organ (Figure 7G). Both *Or85c* and *Or85b* were expressed in a single (and likely the same) bilaterally-symmetric, *Orco*-positive neuron in the dorsal organ in *D. melanogaster* and *D. sechellia* (Figure 7H-J). In *D. melanogaster*, *Or45a* is also expressed in a single neuron, as described previously (Fishilevich et al., 2005). Notably, however, *D. sechellia Or45a* was sometimes detected in two neurons of the dorsal organ (Figure 7I-J). Consistent with the differences in endogenous expression of *Or45a* in these species, *DmelOr45a-Gal4* also almost always labeled a single neuron per dorsal organ, while *DsecOr45a-Gal4* frequently labelled 2-3 neurons (Figure 7K-M). These results suggest that *cis*-regulatory changes in *D. sechellia Or45a* not only underlie the novel adult expression of the receptor in this species, but also the expansion of the expression pattern in larvae.

## Discussion

Neuroethological analysis of animals’ adaptations to ecological niches presents a significant challenge. Studies of species in the field can highlight the complexity of multitrophic interactions of a species with food sources, competitors and predators, but usually lack the ability to probe the neural mechanisms underlying behavioral phenomena. By contrast, laboratory studies have greater experimental accessibility but typically capture only a fraction of the complexity in nature. Here we have sought a “middle-ground” by investigating the interaction of an ecological specialist, *D. sechellia*, at multiple life stages, with its noni host, over the natural ripening process of this fruit. The original description of *D. sechellia*, nearly 50 years ago (Tsacas and Bächli, 1981), already recognized the species’ close – and now generally-believed exclusive – association with noni. This fly has become a classic model for exploring host choice among the myriad possible fruits available in nature (Auer et al., 2021; Jones, 2005). We have extended appreciation of this specialization by revealing a remarkably narrow temporal niche preference: over the many weeks between a noni fruit forming on the *M. citrifolia* tree and its potential consumption by various invertebrate and vertebrate species, our data indicate that *D. sechellia* selects fruit within just a few days of its initial ripening. We discuss the selective advantage of this preference, its mechanistic basis, and the broader insights into the underlying evolutionary processes (Figure 7N).

Our demonstration that both adult and larval *D. sechellia* prefer the noni stage that is most toxic for competitor drosophilids, parasitoids and microbes provides explicit evidence supporting long-held, but largely untested, hypotheses about the fitness benefit of use of this ecological niche. Elimination of competitor drosophilids is likely not only important to preserve a unique food resource for *D. sechellia*, but also beneficial as a reproductive barrier, as this species can naturally interbreed with *D. simulans* (Matute and Ayroles, 2014). The anti-parasitoid and anti-microbial effects of ripe noni appear to also have broader consequences on how this species interacts with these organisms, through reduction or loss of defenses against parasitoid wasps (extending previous observations (Salazar-Jaramillo and Wertheim, 2021)) and pathogenic microbes (O’Malley et al., 2023). Such traits might render *D. sechellia*’s specialization an evolutionary “dead end” (Day et al., 2016), constraining this species to this toxic niche. Finally, one of the more unexpected results of these experiments is the fitness trade-off that *D. sechellia* makes through its preference of ripe noni: at least under the conditions of our assays, ripe noni supports the development of this species less well than overripe stages.

Ripe fruit displays a number of distinct sensory qualities, but we have shown that temporal specialization is guided principally (if not exclusively) by olfactory cues. Previous work identified several olfactory receptors underlying *D. sechellia*’s attraction to, and oviposition on, noni, notably Or22a and Ir75b (Alvarez-Ocana et al., 2023; Auer et al., 2020). However, the odors they recognize are not characteristic of ripe noni, and these receptors are largely dispensable for discriminating ripening stage. Through unbiased chemical profiling of noni across ripening stages and comparative single-cell transcriptomic analysis of the olfactory system, we identified roles for Or45a and Or85c/b in detecting ripe noni-enriched ketones and mediating ripe noni preference. The key contributions of these receptors are further emphasized by the conservation of their requirement for larval ripe noni preference, despite the substantial anatomical and molecular differences in the adult and larval olfactory systems (Vosshall and Stocker, 2007).

While we have been able to identify critical mechanistic elements explaining temporal niche preference of *D. sechellia*, it is important to appreciate that the behavioral and cellular phenotypes are a static snapshot within a continuous process of change on an evolutionary scale. The shift toward use of ripe noni in the *D. sechellia* lineage presumably required coordinated evolution of both toxin resistance and olfactory detection. Noni resistance is a multigenic trait, likely involving cuticular, detoxification and metabolic adaptations (Hungate et al., 2013; Jones, 1998; Lanno et al., 2019; Marconcini et al., 2025; Marconcini et al., 2026). Olfactory detection of ripe fruit is also complex, but here we have been able to provide several insights. First, the two neural pathways required for ripe noni preference in adults have evolved in different ways: Or85c/b neurons retain conserved receptor expression patterns and function in *D. sechellia* (Auer et al., 2020), exhibiting instead an expansion in population size, which affects downstream processing of odor-evoked signals (Takagi et al., 2024). By contrast, Or45a neurons have resulted from derived expression in *D. sechellia* of a receptor gene that was likely only expressed ancestrally in larvae. Second, the novel expression of Or45a in adult Or85a neurons creates a rare example of co-expression of two functional receptors in an OSN population. Little is known about the role of Or85a, but we have shown that it does not contribute to the detection of noni odors in *D. sechellia*. We speculate that this neuronal population represents an intermediate state in olfactory pathway evolution of *D. sechellia*: *cis-*regulatory changes in the *Or45a* promoter enabled co-option of a neuronal pathway by this receptor, coupling noni detection to behavior without the need to generate a completely new sensory pathway. Interestingly, the observed heterogeneity in neuronal spike amplitude of ab2 sensilla subtypes – which likely reflects changes to neuronal morphology (Nava Gonzales et al., 2021; Zhang et al., 2019) – hints at other evolutionary changes in these neuron types, beyond the acquisition of Or45a expression. Third, the rare requirement for two receptors in both adults and larvae underlines the importance of noni discrimination throughout the *D. sechellia* life cycle. As both Or45a and Or85c/b are also expressed in *D. melanogaster* larvae, the neuronal basis underlying the species-specific, noni-stage preference in larvae remains unclear. The atypical expanded Or45a neuronal expression pattern in *D. sechellia* – which we suspect reflects co-option of expression in other Or neuron types rather than additional Or45a neurons *per se* – might provide part of the answer. As this is not a fully penetrant trait, it is possible that distinct central processing of olfactory signals, as described in the larva of another host specialist, *Drosophila erecta* (Roberts et al., 2025), are also relevant. Fourth, full temporal discrimination of noni ripening must involve additional olfactory pathways, as in both adults and larvae, mutants lacking *O45a* and *Or85c/b* (as well as adult *Orco* mutants) still retained aversion to unripe fruit. We suggest that Ir-dependent pathways mediate discrimination of this noni stage.

Together, our work provides unprecedented insights into how chemical changes in an ecological niche can select for molecular and cellular phenotypic divergence in the olfactory system to permit species’ temporal partitioning of a common resource. Such a paradigm represents an exceptional model to understand how the dynamics of environmental selection pressures drive neuronal and behavioral evolution.

## Methods

### Noni fruit sample preparation

*Morinda citrifolia* shrubs were grown under 16:8 h light:dark cycle at 25°C and 50% humidity in the University of Lausanne greenhouses. Unripe fruits are characterized by firmness, green coloration and smaller size, progressively softening and yellowing in the days preceding ripening (Almeida et al., 2019). Fruits typically fall from the tree just before ripening, with full softening occurring in the subsequent few hours. Noni fruits were harvested 1-3 days before full ripening, based on their color (green-yellow) and size. Collected fruits were kept at 30°C and aged for the desired number of days. Fruits of specific ripening stages were mashed with a pestle for 1-2 min to obtain the pulp. As unripe fruits are harder than ripe/overripe fruits, they were ground using a chopper for ∼2 min to reach a pulp-like consistency. All samples were stored at −20°C until use, and freeze-thawed a maximum of two times. For experiments assessing embryonic/larval viability and microbial titers, fresh fruit were used.

### Drosophilid culture

Drosophilids were grown on wheat flour/yeast/fruit juice medium at 25°C under a 12:12 h light:dark cycle. *D. sechellia* stocks were supplemented with noni paste, consisting of Formula 4-24® instant *Drosophila* medium and noni juice (Raab Vitalfood Bio). Wild-type, mutant and transgenic strains are listed in Table S1. *D. melanogaster/D.sechellia* hybrids were generated by crossing 5-10 day old *D. sechellia* males with virgin *D. melanogaster* females (1-3 h post-eclosion); crosses and subsequent embryonic development were maintained at 19°C to maximize survival of hybrids.

### Parasitoid wasp culture

*Leptopilina boulardi* G486 avirulent strain (Russo et al., 1996) and *Leptopilina boulardi* NS1c (Varaldi et al., 2006) were maintained on standard *Drosophila* medium supplemented with honey. To maintain these parasitoid wasp strains, 2-6 day-old mated females were transferred to vials containing second instar *D. melanogaster* Oregon-R larvae and allowed to oviposit. Vials were then kept at 22°C under a 12:12 h light:dark cycle. Around four weeks later, newly emerged adult wasps were collected and staged as described above.

### *Drosophila* mutagenesis and transgenesis

#### D. sechellia Or45a mutant

CRISPR-mediated mutagenesis of *D. sechellia Or45a* (GM21116/LOC6608338) was performed by WellGenetics Inc., adapting previous methodology (Kondo and Ueda, 2013). In brief, oligonucleotides encoding the upstream sgRNA (CCAGCTACTTTGCCGTCCAG[AGG]; PAM in square brackets) and the downstream sgRNA (CTTGCAGGTTATCAGAACGG[CGG]) were cloned separately into a *U6 promoter* plasmid. The 3xP3-RFP cassette, containing a floxed 3xP3-RFP flanked by a 5’ homology arm (−981 nt to −8 nt relative to the *Or45a* ATG) and 3’ homology arm (+1153 nt to +2129 nt relative to the *Or45a* ATG) were cloned into *pUC57-Kan* as a donor template for homology-directed repair. Plasmids encoding the sgRNAs and the repair template were injected into *Dsec.07 pBAC(nos:Cas9,3P3-YFP)* embryos (Auer et al., 2020). An F1 fly expressing the 3xP3-RFP selection marker was crossed to isogenicity, and verified by genomic PCR and Sanger sequencing. The mutant allele, *Or45a^1^*, comprises a 1159-bp deletion of *Or45a* (−7 nt to +1152 nt relative to the ATG), which is replaced by the *3xP3-RFP* cassette.

#### Or45a promoter-Gal4

∼0.7 kb fragments upstream of the *Or45a* start codon were amplified from genomic DNA of *D. melanogaster* Canton-S (using primers (5’-3’) GGCCTGGCAAATGGCTATTTCACTCG and CTTTCAACCGGATTTCGTTGTTGCAC) and *D. sechellia* 07 (using primers GGCCTGGCAAATGGCTATTTCACTCG and CTTTCAACCGGGTTCCGTTGTTGCAC) and inserted upstream of *Gal4* in the *pBPGUw* vector (Addgene #17575) using Gateway cloning. The resultant constructs were integrated into the *D. melanogaster attP2* landing site via phiC31-mediated transgenesis, performed by BestGene Inc.

### Behavior

#### Trap assays

two-choice and multi-choice trap assays were performed following a modified protocol from (Auer et al., 2020). Groups of 25 flies (4- to 7-day-old mated females, unless otherwise indicated) were collected under CO_2_ anesthesia and allowed to recover for ∼24 h in food vials. Flies were starved in empty vials and provided with water-soaked tissue paper for 20-24 h. For each trap (Semadeni #1696, #1511), 250-300 mg of noni pulp were added prior to the assay, and the traps were placed equidistantly in the arena (each trap ∼4 cm from the center) (Semadeni #2962). Flies were briefly anesthetized on ice and introduced into the arena, which was then covered with gauze and kept in the dark – to eliminate visual cues, which are unlikely to contribute to fruit stage discrimination in *D. sechellia* (Alvarez-Ocana et al., 2023) – for 24 h at 25°C and ∼50% humidity. The number of flies in each trap and those remaining outside the traps (no choice) were quantified. For two-choice assays, a preference index was calculated as: (number of flies in the trap “A” - number of flies in the trap “B”)/total number of flies. For multi-choice assays, the percentage of flies in each trap on each substrate was calculated as: (number of flies in each trap/total number of choosing flies) × 100. Arenas in which >25% of flies were found dead were excluded from the analysis. Since trap percentages are compositional (summing to 100% per arena), data were centered log-ratio (CLR) transformed before statistical analysis.

#### Larval chemotaxis assays

multi-choice chemotaxis assays were performed as follows. Second instar larvae were collected by washing them out of the fly food with water; debris and excess water were removed using a fine mesh net. Chemotaxis assays were performed in Petri dishes (Sarstedt #82.1473) covered with 1% agarose (v/v in water). Food patches were prepared by placing Whatman paper disks equidistantly (each disk ∼4 cm from the center) from the center on the agarose layer and adding 70-100 mg of noni pulp of the desired ripening stage on top of each disk. For each experiment, 20-25 larvae were gently collected from the net using a brush and placed in the center of a plate. The plate was covered and placed for 10 min in the dark, at 25°C and ∼70-80% humidity before quantifying the number of larvae on each food patch and those remaining on the agarose surface. The percentage of larvae on each food patch was calculated as: (number of larvae on each patch/total number of choosing larvae) × 100, where choosing larvae refers to animals found on any food patch. Since larval percentages are compositional (summing to 100% per dish), data were centered log-ratio (CLR) transformed before statistical analysis.

#### Multi-choice oviposition assays

groups of 25 4-6-day-old mated females were added to arenas (Semadeni #2962) containing four open tubes (Semadeni #1696) each containing 250-300 mg of noni pulp at different ripening stages. Flies were allowed to lay eggs for 24 h in the dark, at 25°C and ∼50% humidity. Eggs deposited on each substrate were counted after 24 h. For these experiments, both *D. sechellia* and *D. simulans* were reared on standard medium, to exclude the impact of diet on oviposition behavior.

#### No-choice oviposition assays

groups of 10 4-6-day-old mated females were briefly anesthetized with CO_2_ and placed into tubes (Semadeni #1696) containing 250-300 mg of noni pulp. Tubes were covered with a net, and females were allowed to lay eggs for 24 h in the dark, at 25°C and ∼50% humidity. Eggs were counted after 24 h. For these experiments, both *D. sechellia* and *D. simulans* were reared on standard media, to exclude the impact of diets on oviposition behavior.

#### Oviposition retention assays

groups of 10 4-6-day-old mated *Drosophila* females and two males were placed in vials containing standard fly medium. In the test condition, 3 2-5-day-old mated female *Leptopilina boulardi* (strain NS1c) were added; for parallel control conditions, no wasps were added. Vials were kept at 25°C for 24 h, after which flies and wasps were removed and eggs manually counted.

#### Chemotaxis assays using wasp body wash

wasp body wash was extracted as previously described (Davis et al., 2022). Briefly, 70 2-5-day-old mated female wasps were collected in 1.5 ml tubes and stored at −20°C for at least 20 h. For extraction, 350 µl of dichloromethane (DCM; Thermo Scientific #406920010) were added to each wasp tube and vortexed for 2 min. Aliquots of wasp body wash were stored at −20°C until use. For each chemotaxis assay, 15 µl of DCM (as solvent control) or wasp body wash was applied to a Whatman paper disk. Groups of 25 second instar larvae were placed at the center of 1% agarose plates and allowed to choose for 10 min at 25°C in the dark. Rather than counting only larvae located directly on the odor patches, larvae were scored by area: those in the half of the plate containing the control patch and those in the half containing the test patch. The preference index was calculated as follows: (larvae in control area - larvae in test area)/total number of larvae.

### Wasp parasitism

Fifty second-instar drosophilid larvae were placed into a 35 mm dish (Falcon #353001) containing a thin layer of standard fly medium. Five 2-5-day-old mated female wasps (*Leptopilina boulardi* strain G486) were added to the dish and left for 3 h to permit parasitism. Wasps were removed and larvae were transferred to a standard fly medium vial and allowed to develop to adulthood. The following outcomes were scored: parasitized flies (identified by the presence of melanized wasp eggs in the fly abdomen), unparasitized flies, eclosed wasps, and non-emerged animals (wasps or flies).

## Survivorship and viability assays

To assess the survivorship of adult flies and wasps on noni, 2 g of noni pulp of the desired ripening stage were placed in a 35 mm dish (Falcon #353001). Animals were aged 4-6 days post-eclosion, allowed to mate, and briefly anesthetized with CO_2_ immediately before the experiment. Ten mated females were collected and transferred rapidly to dishes containing the pulp. Animals were allowed to recover for 2-3 min before survival monitoring began. Mortality was recorded every 5 min over a 2 h period. For experiments with octanoic acid (Sigma #C2875), 1.5 µl of pure chemical, or diluted in paraffin oil (Fluka 76235, CAS no. 8012-95-1)), were added to the center of the dish.

Viability assays on noni and standard media were performed as follows. *D. sechellia* and *D. simulans* adults were allowed to lay eggs for 20 h on noni juice plates (2.5% agar, 2.5% sucrose, 25% noni juice, mixed in water). Embryos were collected immediately after this period, while first and second/third instar larvae were collected after 48 h and 72 h, respectively (water was added to the plates to prevent desiccation). Embryos and larvae were washed off plates with water using a brush and rinsed to remove any remaining food. For each replicate, 100 embryos or larvae were gently collected using a pipette with cut tips and transferred onto the desired substrate. Viability was monitored daily by counting newly formed pupae and adults.

### Competition assays

To assess the competitive ability of *D. sechellia* relative to *D. simulans*, we performed competition assays on standard media and ripe noni pulp. Groups of 10 4-5-day-old mated females of *D. sechellia* and *D. simulans* were allowed to oviposit either alone or in interspecific competition. For these experiments we used *white* mutant *D. simulans* (*Dsim* 195) to facilitate species identification of the offspring. In single species control tubes, 10 mated females of one species were used. In mixed test tubes, 5 mated females of each species (i.e., 10 total) were added. Females were allowed to lay eggs for 24 h at 25°C and ∼50% humidity, after which they were removed. Tubes were maintained at 25°C, and emerging adults counted. In mixed-species tubes, the two species were distinguished by eye color; the proportion of each species was calculated as the number of emerged adults of that species divided by the total number of emerged adults.

### Bacterial load quantification

Flies were reared on standard medium, ripe (day 0) or late overripe (day 20) noni. Newly-emerged flies were transferred to a fresh tube of the same medium, and colony-forming unit (CFU) analysis was performed as previously described (Montanari et al., 2024). In brief, after 2-3 days, flies were anesthetized and 10 females per condition were collected into Eppendorf tubes containing 600 µl of D-mannitol, MRS, or LB broth and briefly kept on ice. Whole flies were homogenized with a sterile pestle, and two serial 10-fold dilutions (1:10, 1:100; final volume 1 ml) were prepared. For each dilution, 150 µl of whole-fly lysate was plated in triplicate. To recover the major cultivable groups of the *Drosophila* microbiota, homogenates were plated on three media: MRS for lactic acid bacteria (e.g., *Lactobacillus* spp.), D-mannitol agar for acetic acid bacteria (*Acetobacter* spp.), and LB agar for Enterobacteriaceae and other taxa. Plates were incubated at 37°C (MRS) or 30°C (LB and D-mannitol), and colonies were counted after 24 h; overcrowded or confluent plates were excluded from counting. CFU per fly was then calculated as in (Montanari et al., 2024).

### Analysis of noni volatiles

Samples of noni fruits of the desired ripening stage were prepared as described above and stored at −20°C. Three biological replicates per stage were used to assess the consistency of the volatile profile within each stage and to confirm the robustness of our staging. The composition of the volatiles produced by the noni samples was analyzed using solid-phase microextraction coupled with gas chromatography-mass spectrometry (SPME-GC-MS). Samples (1 g/replicate, with three independent biological replicates per ripening stage) were thawed for 3 h before collection of volatiles. The collection of the headspace was carried at 25°C for 30 min, using the SPME fiber assembly (DVB/CAR/PDMS, Supelco®). The front inlet mode was splitless, set up at 240°C, using an Agilent liner (5181-3316). The HP-5ms UI column (Agilent 19091S-413U was fitted in an Agilent 6890N gas chromatograph coupled with MS (5975B). For the column temperature profile, the analysis began at 50°C; this temperature was held for 1 min and gradually increased (3°C/min) to 150°C before holding for 1 min. Subsequently, the temperature was increased (20°C/min) to 260°C and held for 5 min. The MS transfer-line, source and quad were held at 260°C, 230°C and 150°C, respectively.

Raw GC-MS data were exported to AIA format using MSD ChemStation (Agilent Technologies). The exported files were loaded into R (4.1.0) and the XCMS package was used for peak detection and retention time alignment (Smith et al., 2006). In XCMS, the centWave algorithm was used for peak detection using the following parameters: Δm/z of 30 ppm, minimum peak width of 3 s, maximum peak width of 50 s, and signal-to-noise threshold of 20. Retention time correction was performed using the obiwarp function and, for the grouping, an *m*/*z* width of 0.1, base width of 5 and minimum fraction of 0.1 were used. All chromatographic peaks before 50 s and after 1980 s were excluded.

Volatile compounds were identified using the NIST library and matched to standards of the Max-Planck Institute for Chemical Ecology library. GC-MS data were processed as previously described (van den Berg et al., 2006). Briefly, peak intensities of each detected compound were sum-normalized sample-wise, making samples comparable regardless of differences in total volatile emission. Normalized data were then log_10_-transformed to reduce the skewed distribution of the data and subsequently autoscaled to give equal contribution to all detected compounds in our analysis, including minor volatiles that would otherwise be masked by highly abundant compounds.

Principal Component Analysis (PCA) was performed on these “fully processed” data to identify compounds that vary consistently across ripening stages regardless of their absolute abundance. As an alternative approach, PCA was also performed on sum-normalized data only (“minimally processed” data). In this case, the analysis is driven by the most abundant volatiles, which might be ecologically relevant for *D. sechellia* attraction to ripe fruit. Compounds ranking in the top 15 loadings of at least one principal component (PC1 or PC2) in both analytical approaches were considered robust candidate volatiles, as they are both sufficiently abundant and consistently variable across ripening stages.

### Analysis of noni sugars and amino acids

Free amino acids and soluble sugars were extracted from 2.1-3.3 g of fruit tissue samples using 20 ml of methanol followed by centrifugation at 13,000 g for 5 min at 4°C.

Amino acids were quantified with an LC-MS/MS after diluting the methanol extracts 1:10 (v:v) with water containing 10 µg/ml of a mixture of U-^15^N/U-^13^C labeled amino acids (algal amino acids ^13^C, ^15^N, Isotec, Miamisburg, US) and 5 μM of D5-tryptophan (Cambridge Isotope Laboratories, Inc.; Andover, MA). The analysis method was modified from a previous protocol (Crocoll et al., 2016). Chromatography was performed on an Agilent 1260 HPLC system (Agilent Technologies, Boeblingen, Germany). Separation was achieved on a Zorbax Eclipse XDB-C18 column (50×4.6 mm, 1.8 µm; Agilent Technologies, Germany). Formic acid (0.05%) in water and acetonitrile were employed as mobile phases A and B respectively. The elution profile was: 0-1 min, 3% B; 1-2.7 min, 3-100% B; 2.7-3 min 100% B and 3-6 min 3% B. The mobile phase flow rate was 1.1 ml/min. The column temperature was maintained at 25°C. The liquid chromatography was coupled to a QTRAP6500 mass spectrometer (Sciex, Darmstadt, Germany) equipped with a Turbospray ion source operated in positive ionization mode. The ionspray voltage was maintained at 5500 eV. The turbo gas temperature was set at 650°C. Nebulizing gas was set at 60 psi, curtain gas at 35 psi, heating gas at 60 psi and collision gas at “medium”. Multiple reaction monitoring was used to monitor analyte parent ion to product ion (Table S4). Both Q1 and Q3 quadrupoles were maintained at unit resolution. All amino acids were quantified relative to the peak area of the corresponding ^13^C, ^15^N labeled amino acid internal standard, except for asparagine (using aspartate and a response factor of 1.0).

Soluble sugars were analyzed from the fruit extract at 1:10 dilution in water containing 5 µg/ml ^13^C_6_-glucose (Sigma-Aldrich), and 5 µg/ml ^13^C_6_-fructose (Toronto Research Chemicals, Toronto, Canada), by LC-MS/MS as described (Madsen et al., 2015). Targeted analysis used an Agilent 1200 HPLC system coupled to an API 3200 tandem mass spectrometer (AB Sciex, Darmstadt, Germany). The HPLC was equipped with a hydrophilic interaction liquid chromatography (HILIC) column (apHera-NH2 Polymer; Supelco, Bellefonte, PA, USA), and chromatographic separation was performed using water and acetonitrile as mobile phases A and B, respectively, with a flow rate of 1.0 ml/min. The elution profile was: 0-0.5 min, 80% B; 0.5-13 min, 80-55% B; 13-14 min 55-80% B; and 14-18 min 80% B. The mobile phase flow rate was 1.1 ml/min. The column temperature was maintained at 20°C. The mass spectrometer equipped with a Turbospray ion source was operated in the negative ionization mode. The ion spray voltage was maintained at −4200 eV and the turbo gas temperature was set at 500°C. Nebulizing gas was set at 60 psi, curtain gas at 30 psi, heating gas at 60 psi and collision gas at 4 psi. Multiple reaction monitoring was used to monitor analyte precursor ion to product ion (Table S5). Data were acquired using the software Analyst 1.5.1 and quantification was performed using the software MultiQuant 3.0.3 (Sciex, Massachusetts, USA). The concentrations of glucose and fructose were determined relative to the internal standards of ^13^C_6_-glucose and ^13^C_6_-fructose, respectively. The contents of sucrose (Sigma-Aldrich 16104, CAS 57-50-1), trehalose (Sigma-Aldrich T9531, CAS 6138-23-4) and mannitol (sugar alcohol, Carl Roth 30K2.2, CAS 643-01-6) were calculated based on external standard curves.

### Single-nuclear RNA-sequencing of drosophilid antennae

Single-nucleus RNA-seq (snRNA-seq) was performed using 5-day-old mated adult females of *D. melanogaster* CS, *D. simulans* 04 and *D. sechellia* 07. Tissue dissection, nuclear isolation, RNA extraction, library preparation, and sequencing were conducted in parallel across the three species, with three biological replicates per species. Each replicate consisted of 100-200 third antennal segments harvested by snap-freezing animals in a mini-sieve with liquid nitrogen followed by agitation to break off appendages. Single-nucleus suspensions were prepared, and fluorescence-activated nuclear sorting was performed following the Fly Cell Atlas protocol (Li et al., 2022). For each replicate, 15,000-20,000 nuclei were loaded onto a 10x Genomics Chromium Next GEM Chip to target recovery of 10,000 nuclei. Sequencing libraries were generated using the Chromium Single Cell 3ʹ v3.1 dual-index kit, pooled for cluster generation, and sequenced on the Illumina NovaSeq 6000 platform according to 10x Genomics guidelines. Demultiplexing was performed with bcl2fastq2 Conversion Software (v2.20, Illumina). Raw sequencing data were processed using Cell Ranger (v7.1.0) with intronic reads included (--include-introns). Sequence reads from *D. melanogaster*, *D. simulans*, and *D. sechellia* were mapped to their respective reference genomes (*D. melanogaster*: FlyBase release 6.55; *D. simulans*: Prin_Dsim_3.1; *D. sechellia*: ASM438219v2). In total, we obtained transcriptomes of 24,756, 24,763 and 22,046 nuclei for *D. melanogaster*, *D. simulans*, and *D. sechellia*, respectively. Detailed metrics for the snRNA-seq results can be found in Table S6. Downstream analyses were performed in Seurat (v4.4.0). To integrate and cluster cross-species antennal snRNA-seq atlases, we used 13,124 one-to-one-to-one orthologs shared among the three genomes (Lee et al., 2025) as anchors for reciprocal PCA (RPCA)-based integration across all datasets.

Neuronal cell types in the adult antenna were annotated through iterative clustering followed by marker-based assignment. Putative olfactory sensory neuron (OSN) populations were defined based on expression of chemosensory receptor co-receptors (*Orco*, *Ir8a* and *Ir25a*), and the ab1C population was defined by *Gr21a* expression. Among these cell populations, putative doublet clusters that exhibit expression of non-neuronal marker genes, including *Obp19d*, *trol*, *Os-C* and *ct*, were excluded from the analysis. After non-neuronal filtering, OSN populations (co-receptor- and/or *Gr21a*-positive) were subsetted from the full dataset, and dimensional reduction and clustering were re-performed on the OSN-enriched subset. This reclustering yielded many distinct OSN populations characterized by unique or restricted expression of individual olfactory receptors, consistent with known OSN identities in the adult *D. melanogaster* antenna. OSN populations exhibiting expression of multiple Or/Ir genes were further subjected to additional subclustering, and OSN identities were assigned based on receptor gene expression.

### Histology and imaging

Antennae from 5-10-day-old flies (males and females) were collected, and hybridization chain reaction fluorescence *in situ* hybridization (HCR-FISH) on antennal tissue was performed as described (Mermet et al., 2025). The HCR-FISH probes (against target listed in Table S2, all corresponding to *D. melanogaster Or* sequences), amplifiers, and buffers were purchased from Molecular Instruments (HCR^TM^ RNA FISH (v3.0)). For HCR-FISH in the larval dorsal organ, late first-instar or early second-instar larvae (36-48 h after egg-laying) were collected by flooding food vials with 2-3 ml of PBS-T (1×PBSm 0.1% Triton X-100), and rinsing the larvae into a minisieve (mesh-width = 80 µm). Larvae were then rinsed 2-3 times with 1 ml PBS-T and transferred with forceps to a glass dish on ice containing chilled PBS-T, where they were kept until dissection. Larvae were dissected between the segments T3 and A2, and fixed for 1 h in fixation solution (4% paraformaldehyde in 1×PBS, 3% Triton X-100) on ice without rocking. Post fixation, larval tissue was processed identically to antennal tissue as described in (Mermet et al., 2025).

Immunofluorescence stainings were performed on antennal and larval tissues (isolated as described above) essentially as described (Auer et al., 2020), with the rabbit polyclonal anti-GFP (ThermoFisher Scientific A-6455, 1:1000) and AlexaFluor^TM^-488 conjugated goat anti-rabbit IgG (ThermoFisher Scientific A11034, 1:100). In combined HCR-FISH/immunofluorescence stainings, the HCR-FISH protocol was performed first. After the final SSC-T washes of HCR-FISH, samples were washed in 0.2% Triton X-100 in 1×PBS (3 × 5 mins) and blocked for 1 h in blocking solution (5% NGS, 0.2% Triton X-100 in 1x PBS) before incubation with primary antibody and the subsequent steps of the immunofluorescence protocol. Samples were stored in Vectashield at 4°C until imaging. As dorsal-ventral orientation of dissected larval heads was difficult to distinguish, the tissue was mounted between two coverslips to allow imaging of the dorsal organ from either side.

Images were acquired with a Zeiss confocal microscope LSM 710 using a 40× objective, and processed in ImageJ (https://imagej.net/ij/). Neurons were counted manually using the ImageJ MultiPoint tool.

### Electrophysiology

Single sensillum recordings were performed on female flies (5-10-day-old) using tungsten electrodes, essentially as described in (Auer et al., 2020; Benton and Dahanukar, 2023). Sensilla were identified by their morphology, characteristic location, and responses to diagnostic ligands as listed in the DoOR database (Munch and Galizia, 2016). Odor cartridges were prepared as described (Auer et al., 2020; Benton and Dahanukar, 2023). Odors were diluted in paraffin oil (Fluka 76235, CAS 8012-95-1), except ethyl-*3*-hydroxybutyrate, which was diluted in water. Odor stimuli were delivered to the fly for 1 s, and responses were calculated as the number of spikes in a 500 ms window during odor delivery minus the baseline spike count in a 500 ms pre-stimulus window, and multiplied by 2 to obtain spikes/s. For each sensillum, odor responses were corrected by subtracting the corresponding solvent response to obtain solvent-corrected spikes/s. Each odor cartridge was used for a maximum of five trials. Odors used in this study are listed in Table S3. For cross-species comparisons of ab2 OSN responses, we focused only on the *D. sechellia* ab2 sensilla with two distinct spike amplitudes.

For measuring neuronal responses to noni (Figure 6A,D), fruit pulp was prepared as described above. Following pilot tests, we used as stimuli (in the standard odor cartridges) either 50 or 10 mg of fruit for ab2 and ab3 sensilla recordings, respectively. In ab3 sensilla, Or85b/c neurons were sensitive to noni volatiles, leading us to use the lowest airflow for odor pulse stimulation (“#” on Syntech CS-55) to further reduce odor concentration. Here, each odor cartridge was used for a maximum of one trial.

## Statistical analysis

Statistical analyses were performed in GraphPad Prism (version 11) and R (version 4.5.1). For each experiment, *n* indicates the number of biological replicates and is reported, together with the statistical test used, in the corresponding figure legend. Normality tests were run to assess data distribution, and parametric or non-parametric tests were applied accordingly; post-hoc tests were used to correct multiple comparisons and are reported in the figure legend. Significance was set at α = 0.05.

## Acknowledgements

We thank Michèle Crozatier, Maria Cristina Gambetta, Bruno Lemaitre, Julien Varaldi and Leslie Vosshall for reagents, Blaise Tissot-Dit-Sanfin for cultivation of the *M. citrifolia* plants, and Bill Hansson and Jonathan Gershenzon for support in the chemical analyses. We acknowledge the Bloomington *Drosophila* Stock Center (NIH P40OD018537) for *D. melanogaster* stocks and the Developmental Studies Hybridoma Bank for antibodies. We are grateful to members of the Benton laboratory for discussions and comments on the manuscript. A.M. was supported by an EMBO Long-term Fellowship (ALTF 447-2022). M.R., R.D.M.-S. and M.K. were supported by the Max Planck Society. R.D.M.-S. was supported by the German Academic Exchange Service (DAAD 57440921). D.L. is supported by grants from the National Research Foundation of Korea (NRF), funded by the Ministry of Science and ICT (MSIT) (RS-2023-00211007 and RS-2024-00411768, Bio & Medical Technology Development Program). Research in R.B.’s laboratory was supported by the University of Lausanne, an ERC Advanced Grant (833548) and the Swiss National Science Foundation (310030_219185).

## Author contributions

A.M. conceived the project, and designed and performed all behavioral, viability and bacterial analyses, as well as the data analysis of noni volatiles and RNA FISH validation of *Or45a* mutants. A.S.B. performed all other histological experiments, electrophysiological analyses and generated the *Or45a* promoter transgenes. R.D.M.-S. performed the SPME-GC-MS analyses, supervised by M.K. L.A. performed behavioral, viability and bacterial analyses, and S.C. performed viability analyses, supervised by A.M. M.R. carried out analysis of amino acids and sugars. M.M. assisted with statistical analyses of behavioral data. D.L. designed, performed and analyzed the single-nucleus RNA-seq experiments. R.B. conceived, managed and supervised the project. A.M., A.S.B. and R.B. wrote the paper, with contributions from D.L., R.D.M.-S. and M.R., and feedback from all authors.

## Supplementary Figure legends

**Figure S1. Additional analyses of noni stage ripening preference.** (A) Left: behavioral responses of *Dmel* CS in multi-choice trap assays with four ripening stages. Box plots show the percentage of individuals trapped on each substrate. Right: radar plot comparing mean percentages of trapped flies per stage for *Dsec* 07 (control) and *Dmel* CS (d, day of ripeness). *Dmel* CS (box and radar plots), *n* = 22; *Dsec 07* control (radar plot only), *n* = 17. (B) Behavioral responses of *Dsec* 07 (starved mated females, as control for all other conditions) in multi-choice trap assays with four ripening stages. Box plot shows the percentage of individuals trapped on each substrate; *n* = 24. (C,) Left: behavioral response of fed mated females (C) or starved virgin females (D) in multi-choice trap assays with four ripening stages. Box plots show the percentage of individuals trapped on each substrate. Right: radar plots comparing mean percentages of trapped flies per stage between *Dsec* 07 (starved mated females, as control) and *Dsec* 07 at different internal states (d, day of ripeness). (C) *Dsec* 07 fed mated females (box and radar plots), *n* = 19; control (radar plot only), *n* = 24. (D) *Dsec* 07 starved virgin females (box and radar plots), *n* = 15; control (radar plot only), *n* = 19. (E) Behavioral responses of *Dsec* 07 and *Dsec* 28 in multi-choice trap assays with four noni ripening stages. For these experiments, fruits were staged once they were ripe (day 0) for 1, 2 or 3 days. Box plots show the percentage of individuals trapped on each substrate; *n* = 12 (*Dsec* 07)*, n* = 7 (*Dsec* 28). (F) Behavioral responses of *Dsec* 07 and *Dsec* 28 in multi-choice trap assays testing preferences across ripening stages, excluding the ripe (day 0) stage. Box plots show the percentage of individuals trapped on each substrate; *n* = 21 (*Dsec* 07), *n* = 14 (*Dsec* 28). For all the plots, n represents the number of arenas. All box plots show the median and the first and third quartiles, with points representing biological replicates. In (A-F), statistical analyses were performed on CLR-transformed values. Repeated measures one-way ANOVA with Tukey’s post-hoc test was performed for within-genotype comparisons with significant differences indicated by different letters (ɑ = 0.05). Mixed-design ANOVA with Genotype × Stage interaction f was used for genotype comparisons (radar plots; *p* value for the difference in preference between genotypes is shown).

**Figure S2.** Parasitoid wasp assays and octanoic acid toxicity. (A) Survivorship of *D. sechellia* (*Dsec* 07) and *L. boulardi* (strain G486) adult females on octanoic acid (OA) at different concentrations. Paraffin oil is the solvent control. Survival curves represent pooled data from all experiments; *n* = 10-20 animals per species. (B) Left: schematic of the oviposition retention assay. Five mated *Drosophila* females and two males were placed in vials for 24 h (control condition). In the test condition, three mated *L. boulardi* females were also added. Right: oviposition retention assays using the indicated species/strains, in the presence or absence of *L. boulardi* (strain NS1c); *n* = 12-18 per species and strain. (C) Left: schematic of the two-choice chemotaxis assays using *L. boulardi* (strain G486) wasp body wash and solvent control (dichloromethane). Right: behavioral responses of *D. melanogaster* (strain *Dmel* OR) and *Dsec* 07; *n* = 9-10. (D) Left: Schematic of the parasitism assay using *L. boulardi* (strain G486) and second instar *Drosophila* larvae. After 3 h, larvae were transferred to new vials and allowed to develop to adulthood. Right: Results of parasitism assays using *Dmel* OR, *Dsec* 07, and *Dsec* 28 larvae. Raw counts of unparasitized flies, parasitized flies, emerged wasps, and non-emerged animals are shown; *n* = 3 per species and strain. In (B) and (C), box plots show the median and the first and third quartiles, with points representing biological replicates (*n*). In (A), Log-rank tests with Holm correction. \*\*\**p* < 0.001. In (B), Mann-Whitney test. \*\**p* < 0.01, \*\*\**p* < 0.001. In (C), unpaired t-test. \**p* < 0.05.

**Figure S3.** PC1 and PC2 loadings of all detected noni volatiles. (A) PC loadings for fully processed data. Left: PC1 loadings of all detected volatile compounds. Right: PC2 loadings of all detected volatile compounds. (B) PC loadings for minimally processed data. Left: PC1 loadings of all detected volatile compounds. Right: PC2 loadings of all detected volatile compounds. (A-B) points represent the loading value of each compound. The top 15 compounds by absolute loading value are highlighted in red (positive loadings) or blue (negative loadings). See also Figure 3E.

**Figure S4.** Analysis of sugars and free amino acids across noni ripening stages. (A) Principal Component Analysis (PCA) score plot showing the separation of fruit samples by ripening stage based on their sugars profile. Prior to sugar analysis, peak intensities were normalized, log-transformed and autoscaled. (B) Hierarchical cluster analysis dendrogram of the 12 fruit samples, computed on autoscaled data using Euclidean distance and Ward’s linkage method. The heatmap below shows the relative abundance of detected sugars. Color scale represents z-scores calculated per compound across all samples. (C) PCA score plot showing the separation of fruit samples by ripening stage based on their free amino acids profile. Prior to amino acid analysis, peak intensities were normalized, log-transformed and autoscaled. (D) Hierarchical cluster analysis dendrogram of the 12 fruit samples, computed on autoscaled data using Euclidean distance and Ward’s linkage method. The heatmap below shows the relative abundance of detected free amino acids across ripening stages. Color scale represents z-scores calculated per compound across all samples.

**Figure S5.** Comparative analysis of Ir subsystem OSN populations and cluster analysis of ab1 and ab2 OSNs. (A) tSNE plots of *D. melanogaster*, *D. simulans* and *D. sechellia* antennal Ir subsystem neurons from an integrated dataset after RPCA integration. In the bottom right plot, all cells from the three species are merged. (B) tSNE plot of Ir subsystem neurons from the merged dataset in which cluster identity (neuron type) is annotated. The unassigned neuron types might correspond to additional sacculus neurons, but we lack sufficient information to confidently make such assignments. For example, the two unannotated clusters (here and in (A)) are likely to correspond to sacculus Ir68a neurons, although transcripts of this lowly-expressed receptor (Knecht et al., 2017) were not detected in our dataset. (C) Bar plot illustrating Ir subsystem neuron type frequency comparisons across species. Each point corresponds to one of the three biological replicates. (D) tSNE plots of ab1 and ab2 OSNs from the *D. melanogaster* and *D. sechellia* datasets (reclustered separately), with annotations of OSN types. (E) tSNE plots from (D) highlighting expression of the indicated *Or* genes. Color intensity represents relative SCT-normalized expression of the indicated gene.

**Figure S6.** Electrophysiological characterization of typical and atypical ab2 sensilla in *D. sechellia*, and generation of an *Or45a* mutant. (A-D) Left: representative traces of electrophysiological responses to the indicated odors of *D. sechellia*’s “typical” ab2 sensilla (with two distinct spike amplitudes) and “atypical” ab2 sensilla (which have two neurons of similar spike amplitude; note these neurons can sometimes be distinguished after odor stimulation, when the responding neuron spike train “pinches” slightly, as indicated in (B)); black horizontal bar, odor stimulus (1 s). Right: quantification of dose-responses to the indicated ligand of ab2 sensilla (i.e., summed response from both A and B neurons); *n* = 4-5 sensilla per odor concentration. (E) Schematic of *D. sechellia Or45a* mutant allele generation via CRISPR/Cas9-mediated gene replacement. Data in (A-D) are represented as mean ± SD; Shapiro-Wilk test for normal distribution, followed by unpaired two-tailed Student’s t-test comparison between neuronal responses at each concentration; no significant differences between responses in typical vs atypical sensilla types for odorants at any concentration.

**Figure S7. Central projections of neurons expressing *Or45a* promoter reporters.** (A-B) Representative immunofluorescence for GFP and nc82 on *D. melanogaster* brains of animals expressing *UAS-mCD8:GFP* under the control of *DmelOr45a-Gal4* (A) and *DsecOr45a-Gal4* (B). *DmelOr45a-Gal4* drives only ectopic GFP expression in neurons projecting to the VM2 glomerulus. *DsecOr45a-Gal4* drives expression in neurons projecting to DM5 (*Or85a*), likely representing “real” activity of the *DsecOr45a* promoter sequence), as well as the same ectopic expression in VM2 neurons. Scale bar, 25 µm.

## Supplementary Tables

**Table S1:**
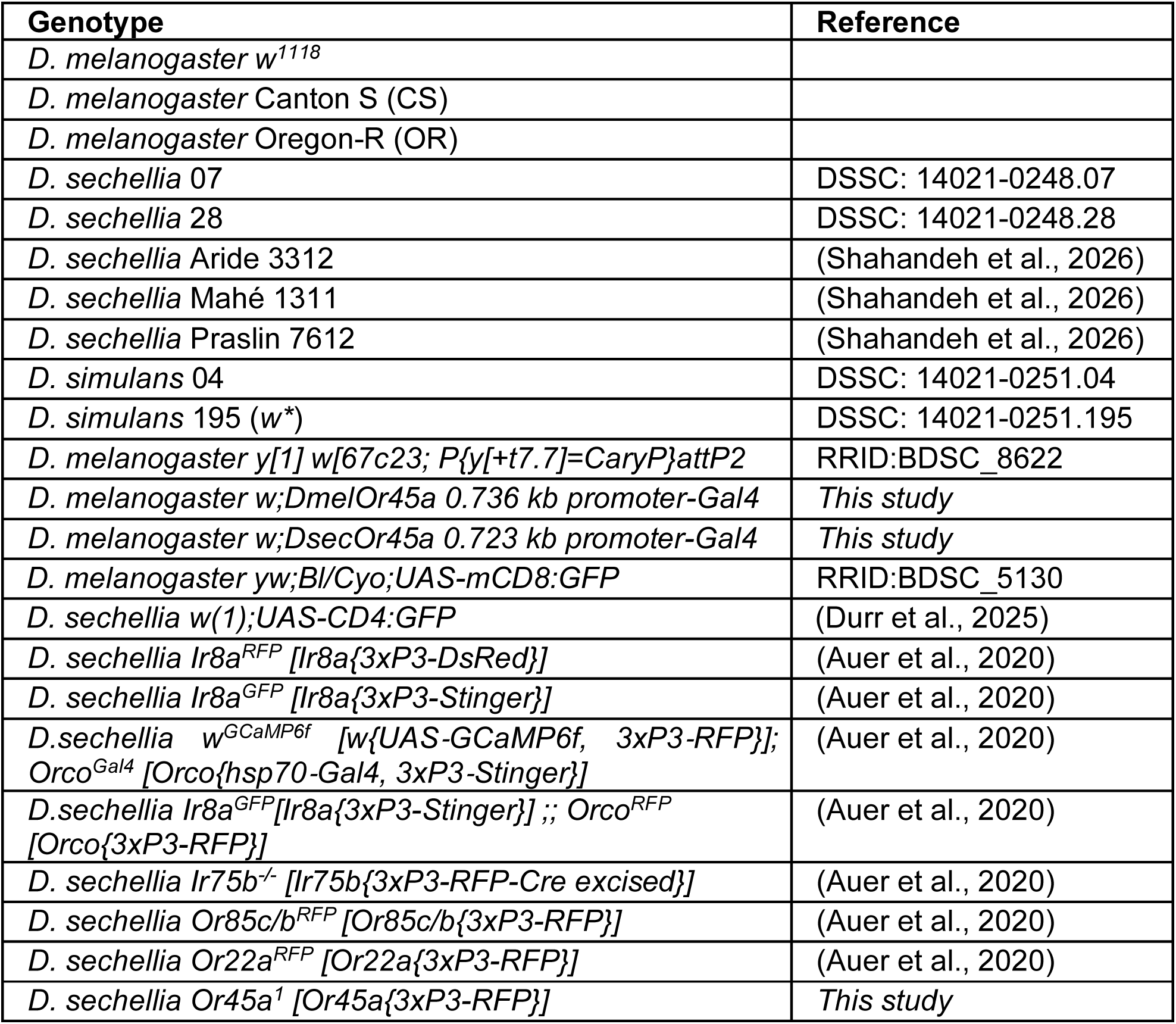
Drosophilid strains.

**Table S2:** HCR RNA FISH probe target sequences.

| Gene | Target sequence |
| --- | --- |
| *Or7a* | NM_078526.1 |
| *Or45a* | NM_078942.4 |
| *Or56a* | NM_079072.2 |
| *Or59b* | NM_079098.2 |
| *Or85a* | NM_079553.1 |
| *Or85b* | NM_079555.2 |
| *Or85c* | NM_079556.3 |
| *Orco* | NM_079511.5 |

**Table S3:** Odors.

| Odor name | Cas | Source |
| --- | --- | --- |
| methyl acetate | 79-20-9 | Sigma Aldrich 296996 |
| ethyl 3-hydroxybutyrate | 5405-41-4 | Sigma Aldrich E30603 |
| 2-heptanone | 110-43-0 | Sigma Aldrich 537683 |
| 2-nonanone | 821-55-6 | Fluka 52000 |

**Table S4:** Details of analysis of amino acids by LC-MS/MS [HPLC 1260 (Agilent Technologies)-QTRAP6500 (AB SCIEX)] in positive ionization mode.

| Compound | Q1 | Q3 | RT<br>(min) | Internal<br>standard | IS<br>Q1 | IS<br>Q3 | DP | CE |
| --- | --- | --- | --- | --- | --- | --- | --- | --- |
| Ala | 90.1 | 44.1 | 0.5 | <sup>13</sup>C,<sup>15</sup>N-Ala | 94.1 | 47.1 | 20 | 17 |
| Ser | 106.0 | 60.1 | 0.5 | <sup>13</sup>C,<sup>15</sup>N-Ser | 110.0 | 63.1 | 20 | 15 |
| Pro | 116.1 | 70 | 0.5 | <sup>13</sup>C,<sup>15</sup>N-Pro | 122.1 | 75.0 | 20 | 19 |
| Val | 118.1 | 72.2 | 0.5 | <sup>13</sup>C,<sup>15</sup>N-Val | 124.1 | 77.2 | 20 | 13 |
| Thr | 120.1 | 74.2 | 0.5 | <sup>13</sup>C,<sup>15</sup>N-Thr | 125.1 | 78.2 | 20 | 13 |
| Ile | 132.2 | 86.1 | 1.1 | <sup>13</sup>C,<sup>15</sup>N-Ile | 139.2 | 92.1 | 20 | 13 |
| Leu | 132.2 | 86.1 | 1.3 | <sup>13</sup>C,<sup>15</sup>N-Leu | 139.2 | 92.1 | 20 | 13 |
| Asp | 134.1 | 74.1 | 0.5 | <sup>13</sup>C,<sup>15</sup>N-Asp | 139.1 | 77.1 | 20 | 19 |
| Glu | 148.1 | 102.1 | 0.5 | <sup>13</sup>C,<sup>15</sup>N-Glu | 154.1 | 107.1 | 20 | 15 |
| Met | 150.2 | 104.1 | 0.7 | <sup>13</sup>C,<sup>15</sup>N-Met | 156.2 | 109.1 | 20 | 13 |
| His | 156.2 | 110.1 | 0.4 | <sup>13</sup>C,<sup>15</sup>N-His | 165.2 | 118.1 | 20 | 17 |
| Phe | 166.2 | 120.2 | 2.6 | <sup>13</sup>C,<sup>15</sup>N-Phe | 176.2 | 129.2 | 20 | 17 |
| Arg | 175.1 | 70.1 | 0.4 | <sup>13</sup>C,<sup>15</sup>N-Arg | 185.1 | 75.1 | 20 | 31 |
| Tyr | 182.1 | 136.2 | 1.4 | <sup>13</sup>C,<sup>15</sup>N-Tyr | 192.1 | 145.2 | 20 | 17 |
| Asn | 133.1 | 74.1 | 0.5 | <sup>13</sup>C,<sup>15</sup>N-Asp |  |  | 20 | 21 |
| Gln | 147.1 | 130 | 0.5 | <sup>13</sup>C,<sup>15</sup>N-Gln | 154.1 | 136.0 | 20 | 13 |
| Trp | 205.2 | 188.1 | 3.2 | D5-Trp | 210.0 | 193.0 | 20 | 13 |

**Table S5:**
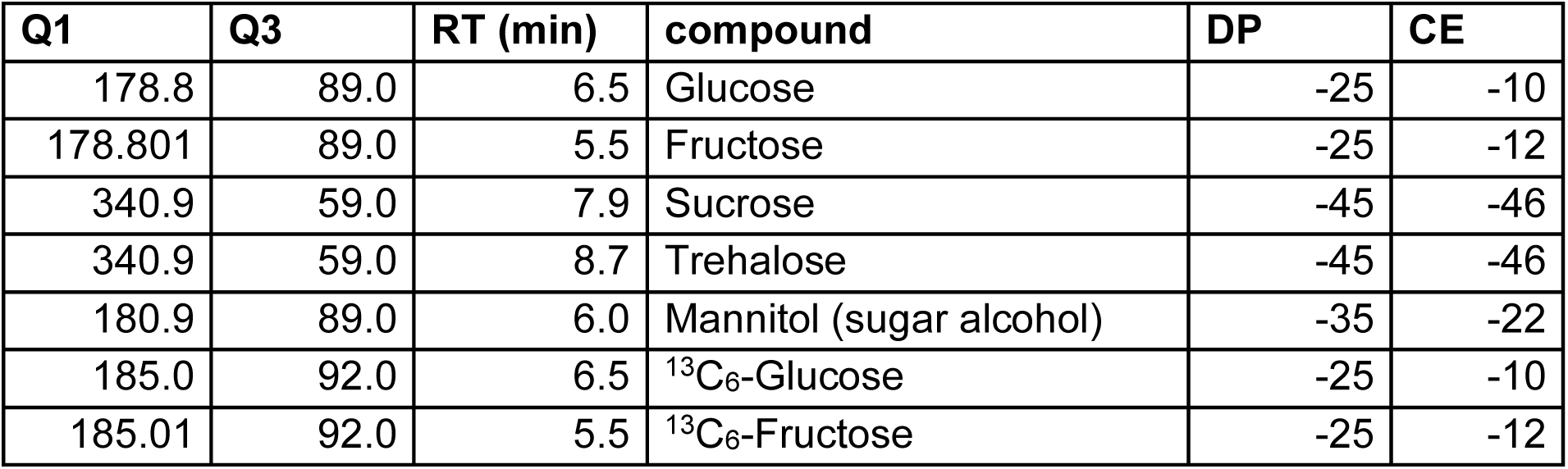
Details of analysis of sugars by LC-MS/MS [HPLC 1200 (Agilent Technologies)-API3200 (AB SCIEX)] in negative ionization mode.

| Q1 | Q3 | RT (min) | compound | DP | CE |
| --- | --- | --- | --- | --- | --- |
| 178.8 | 89.0 | 6.5 | Glucose | -25 | -10 |
| 178.801 | 89.0 | 5.5 | Fructose | -25 | -12 |
| 340.9 | 59.0 | 7.9 | Sucrose | -45 | -46 |
| 340.9 | 59.0 | 8.7 | Trehalose | -45 | -46 |
| 180.9 | 89.0 | 6.0 | Mannitol (sugar alcohol) | -35 | -22 |
| 185.0 | 92.0 | 6.5 | <sup>13</sup>C<sub>6</sub>-Glucose | -25 | -10 |
| 185.01 | 92.0 | 5.5 | <sup>13</sup>C<sub>6</sub>-Fructose | -25 | -12 |

**Table S6:** Metrics for the snRNA-seq antennal atlases.

| Species | Replicate | Estimated Number of Cells | Mean Reads per Cell | Median Genes per Cell | Total Genes Detected | Median UMI Counts per Cell |
| --- | --- | --- | --- | --- | --- | --- |
| *D. melanogaster* | 1 | 5159 | 25494 | 643 | 11855 | 1151 |
|  | 2 | 8956 | 23389 | 628 | 12805 | 1188 |
|  | 3 | 10641 | 21828 | 726 | 12936 | 1456 |
| *D. simulans* | 1 | 3558 | 25937 | 549 | 11130 | 930 |
|  | 2 | 9508 | 23948 | 531 | 12069 | 890 |
|  | 3 | 11697 | 29516 | 595 | 12544 | 1043 |
| *D. sechellia* | 1 | 3184 | 38941 | 589 | 11313 | 1067 |
|  | 2 | 8612 | 24455 | 491 | 12188 | 818 |
|  | 3 | 10250 | 28319 | 584 | 12446 | 1069 |

**Table S7:** Noni odors identified by GC-MS. *Provided as a separate Excel file*

