## Supplementary figures and images for "Temporal niche partitioning through olfactory cell type evolution"

### Figures S1-S7

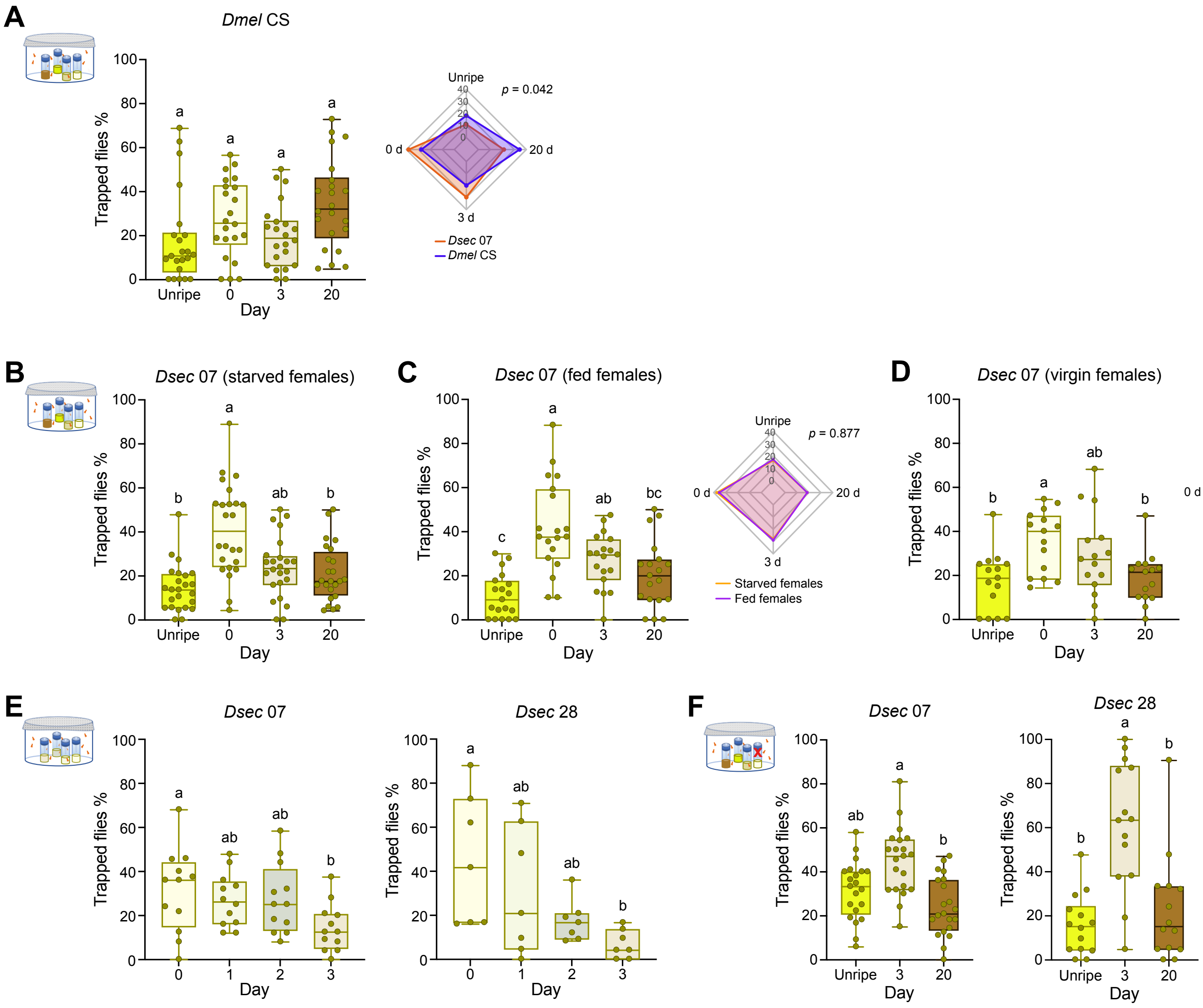

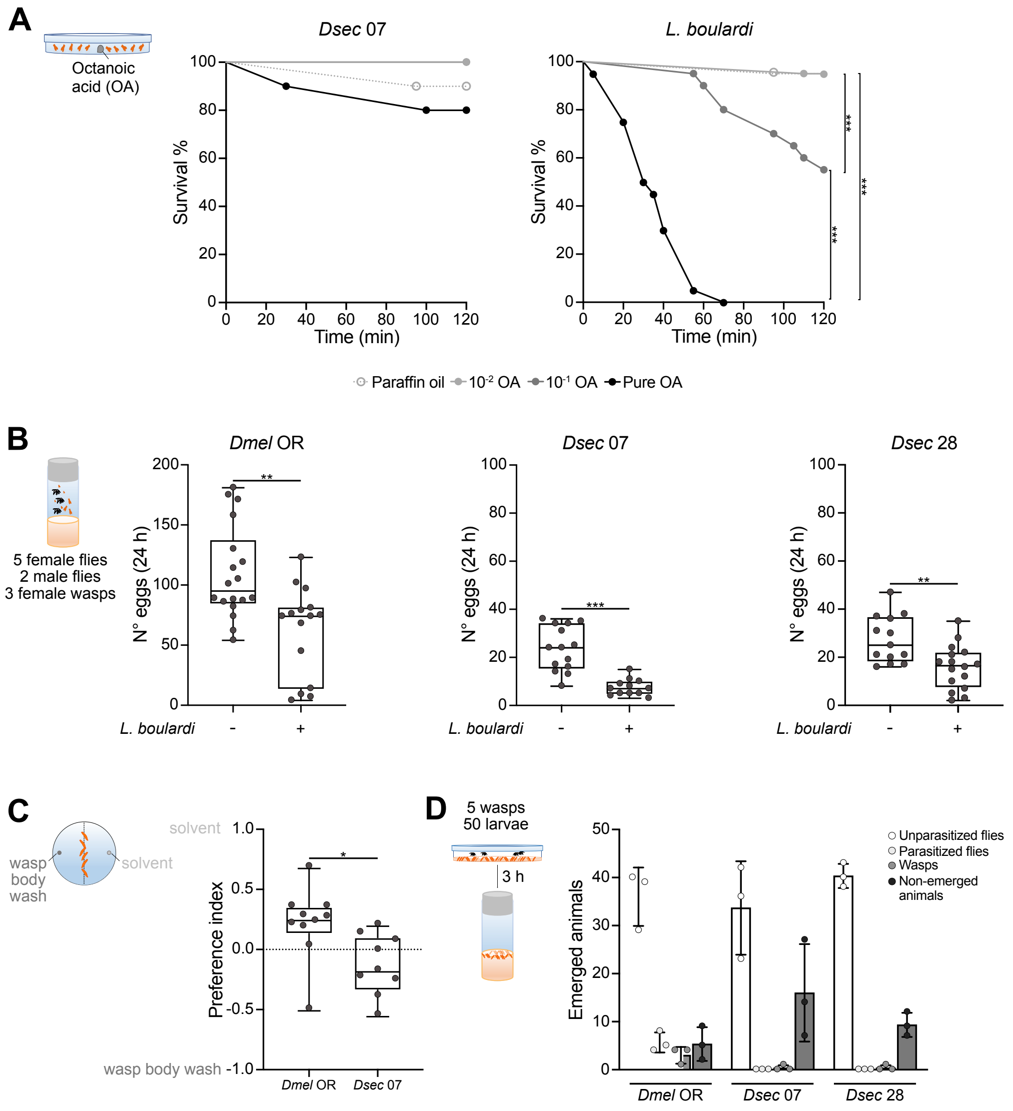

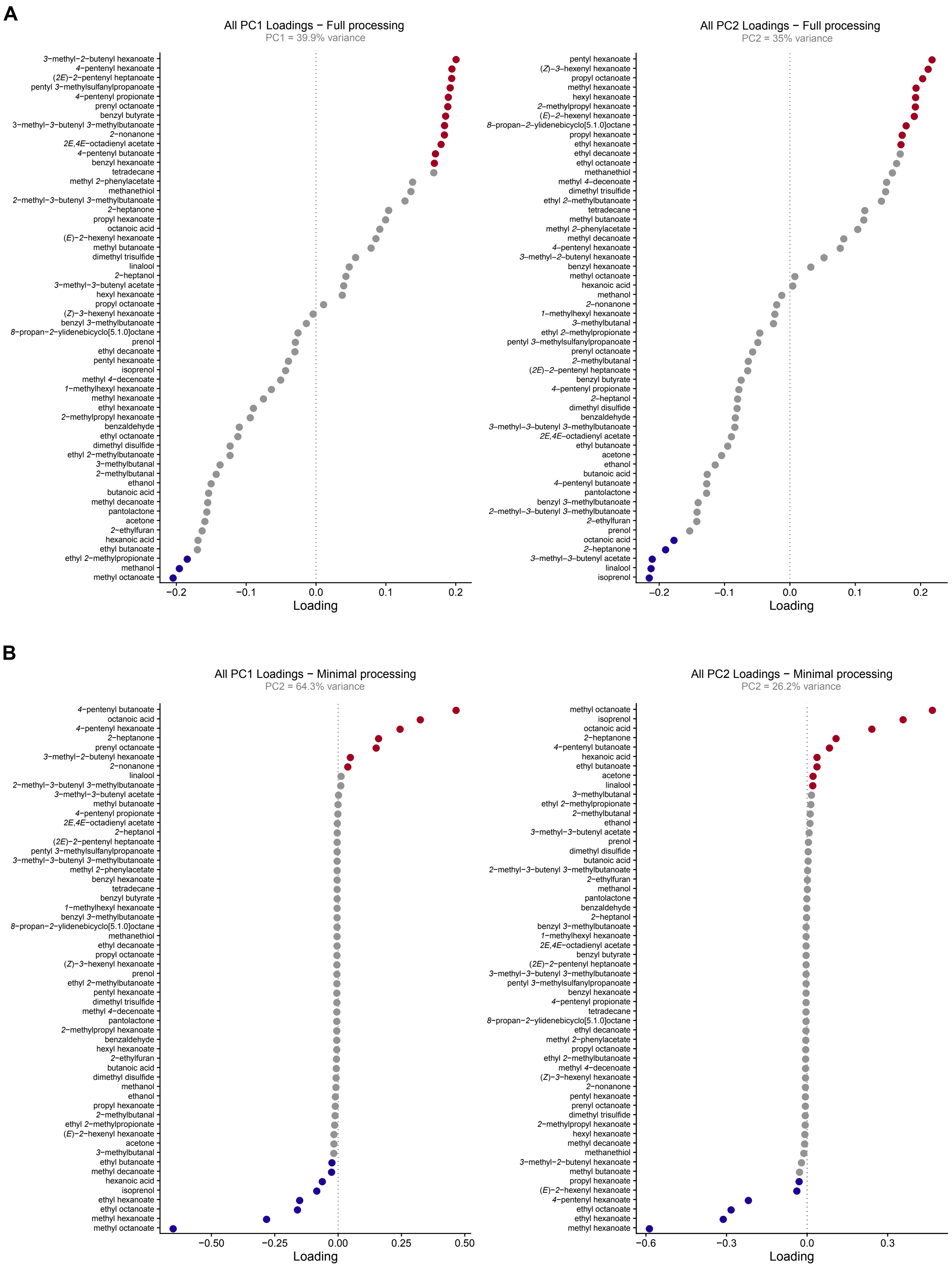

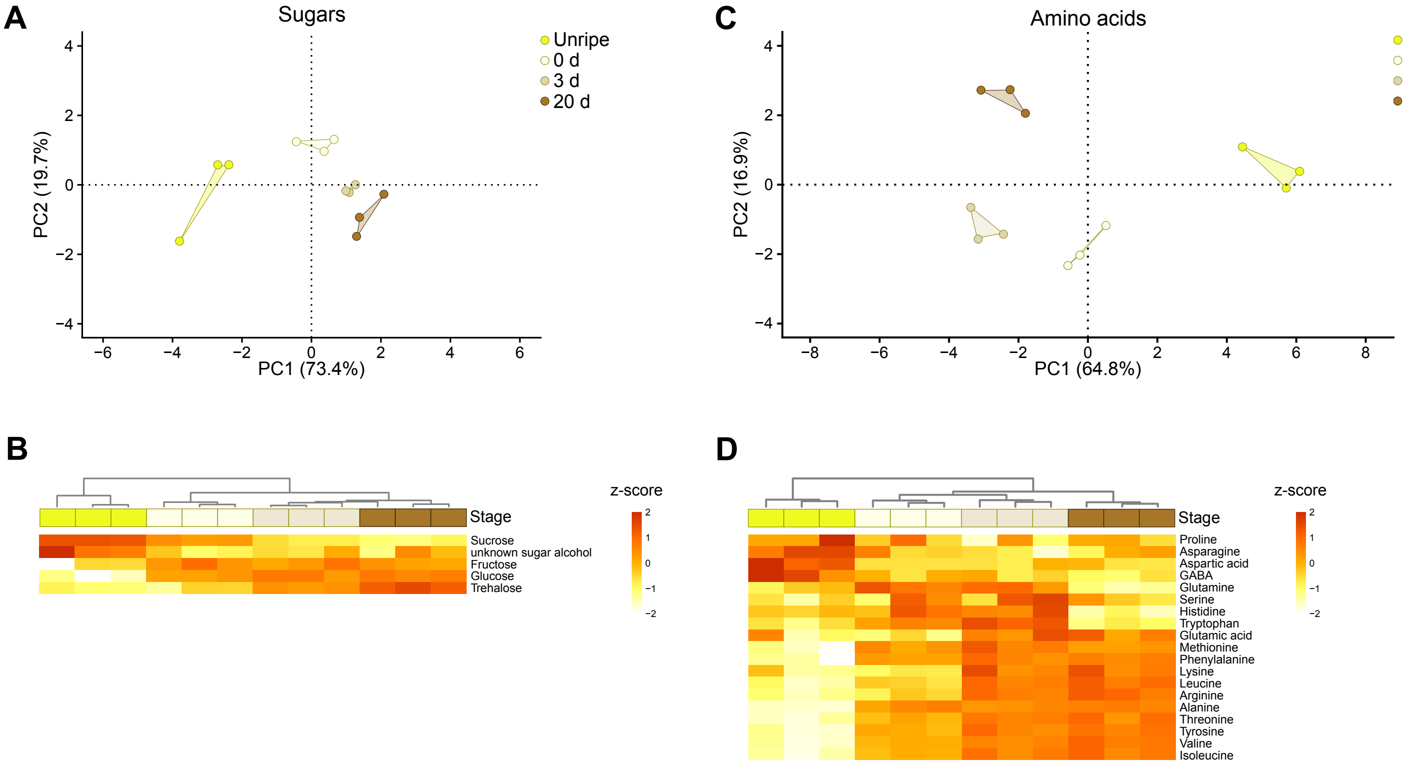

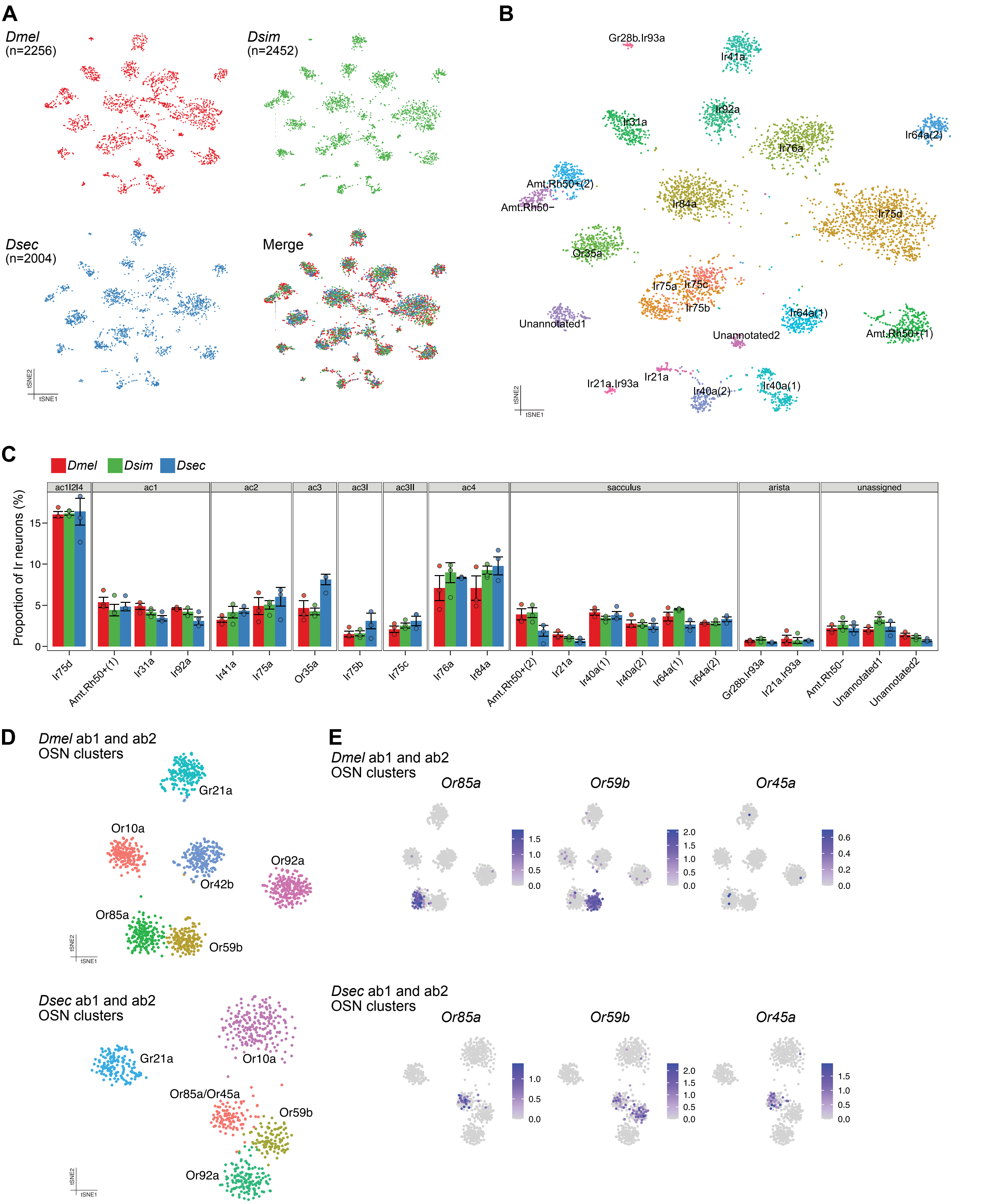

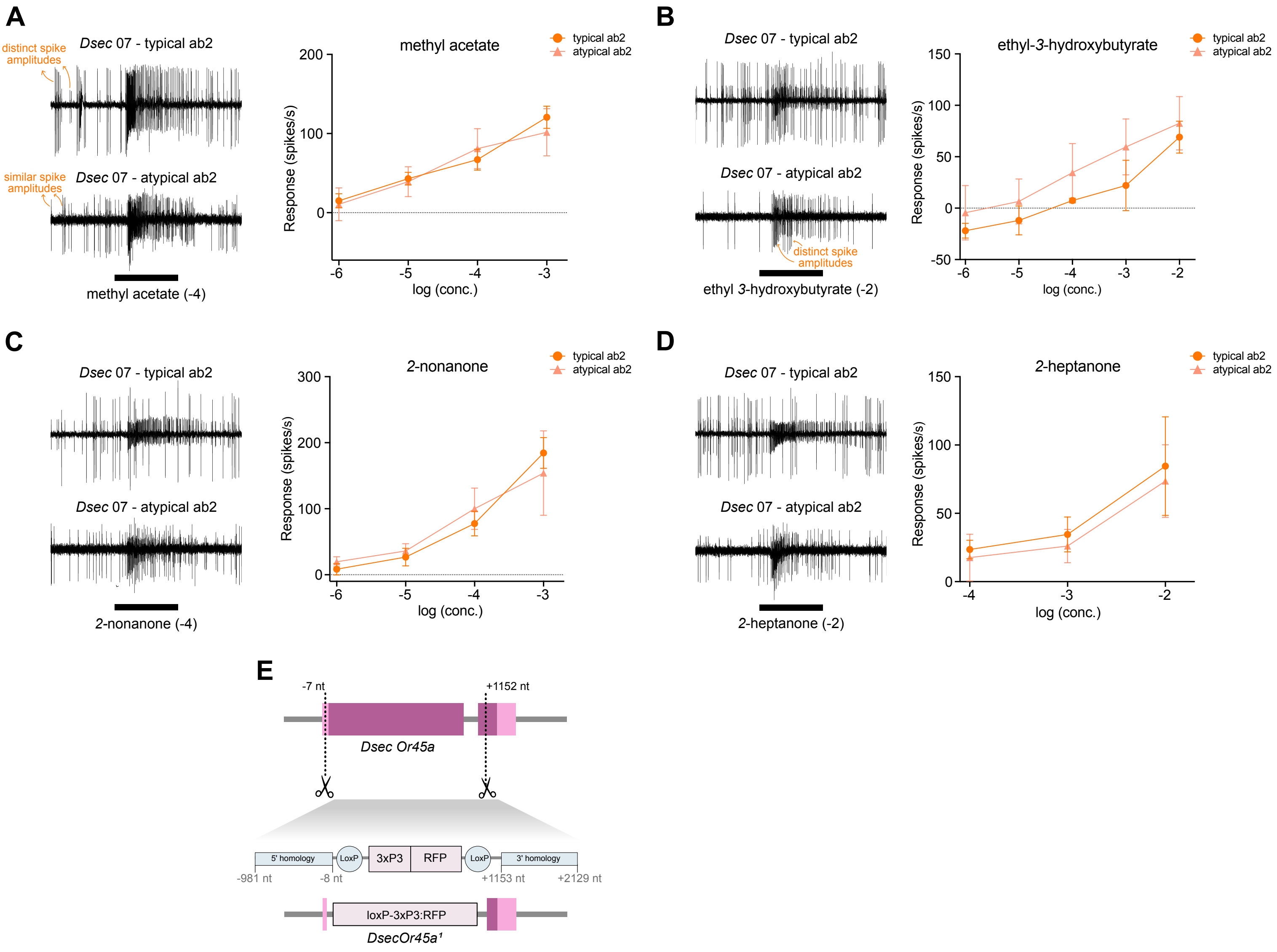

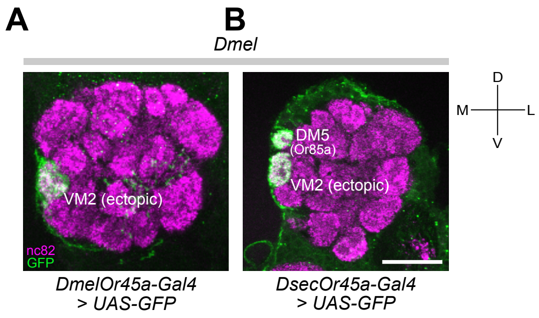
